# Growth dynamics and protein-expression of *Escherichia coli* serotypes O26:H11, O111:H8 and O145:NM in the bovine rumen

**DOI:** 10.1101/2024.11.06.622239

**Authors:** Indira T. Kudva, Erika N. Biernbaum, Julian M. Trachsel

## Abstract

To adapt to the ruminal environment, Shiga toxin-producing *Escherichia coli* (STEC) O157:H7 (O157) expresses proteins involved in survival rather than virulence. Additionally, STEC O157 strains exhibit distinct *in vitro* but shared *in vivo* survival patterns in rumen fluids that sets them apart from non-STEC, commensal *E. coli*. To determine if similar responses would be observed with other STEC, we evaluated three non-O157 serotypes, O26:H11, O111:H8 and O145:NM, along with a non-STEC *E. coli*, under the growth conditions used for STEC O157: (i) anaerobically, *in vitro*, in rumen fluid from cattle on a lactation (low fiber, high protein) or maintenance (high fiber, low protein) diet, at 39°C for 48 hr and (ii) *in vivo* for 48 hr within the rumen of cattle on the same diets using a non-terminal, rumen-fistulated animal model that allows for introduction of bacteria without ruminal contamination. On the lactation diet, the ruminal pH was acidic ranging from 5.2 - 6.0 and the total volatile fatty acids (VFA) concentration, 141 - 230 μM/ml. On the maintenance diet, the ruminal fluid pH was close to neutral ranging from 6.0 - 7.0, with a total VFA of 87 - 190 μM/ml. Unlike STEC O157, the three non-O157 serotypes demonstrated survival patterns similar to each other and the control non-STEC *E. coli.* A greater reduction in viable STEC counts was observed *in vitro* in rumen fluid from cattle fed the lactation diet than *in vivo*, corroborating previous reports that *in vitro* conditions cannot mimic those observed *in vivo*. Like STEC O157, the non-O157 serotypes mainly expressed proteins supporting ruminal adaptation although not all proteins matched those expressed by STEC O157, and included the virulence protein, intimin. Explorative studies such as this could provide insights into common conditions/ targets that may have application in broader STEC control in cattle.

**Importance of this study:** This study demonstrates that non-O157 serotypes O26:H11, O111:H8, O145:NM have similar survival and protein expression patterns in rumen fluid with variations being influenced primarily by rumen fluid composition associated with diet. Unlike STEC O157, under *in vivo* conditions, the growth dynamics of the non-O157 serotypes were comparable to that of non-STEC, commensal *E. coli*. Hence, exploring bacterial protein expression within the host is critical in discerning therapeutic targets, unique to/shared between STEC, for broad control strategies. In addition, this study further validates the value of using a non-terminal animal model for rumen studies that reduces number of animals used for an experiment.

## INTRODUCTION

Cattle are asymptomatic reservoirs of Shiga toxin-producing *Escherichia coli* (STEC), which can cause a range of illnesses in humans, from diarrhea to hemorrhagic colitis (HC) and hemolytic uremic syndrome (HUS) [1, 2]. STEC reported in the top six of 15 leading foodborne pathogens cause a cumulative illness-related economic burden of $17.6 billion in the United States [3], and are also one of four foodborne pathogens most frequently contaminating USDA- Food Safety and Inspection Service (FSIS) regulated products [4]. Worldwide, STEC cause 1.2 million illnesses and 128 deaths each year, according to a World Health Organization risk assessment report [5]. Human illness follows the seasonal STEC-shedding patterns observed in cattle, peaking during the summer months [6–8] *E. coli* O157:H7 (O157) is the most well-studied STEC serotype causing 36% of cases in the United States and 2.8 million cases worldwide [9–11]. However, there are non-O157 STEC serotypes that are also associated with human illness, first documented in the United States in 1994 [12]. In 2012, the USDA-FSIS announced the “Big Six” non-O157 STEC serogroups (O26, O45, O103, O111, O121, and O145) that are ‘commonly implicated’ and isolated from domestic or imported beef and cattle [13]. The CDC estimates that, as a group, the “Big Six” outnumber O157 in human illness cases and it is likely that illnesses attributed to non-O157 STEC have been underestimated due to traditional testing geared towards O157 detection [14].

Increased reporting of non-O157 STEC cases (isolated and outbreaks) in the United States started between years 2004 – 2014, likely due to targeted diagnostics, with the top STEC serogroups including O26, O103 and O111 [12]. The STEC serogroup O26 was most commonly associated with human illness only after O157 [14, 15], with O26:H11 being most prevalent from years 1995-2014 [12]. Globally, clinical cases with serogroup O26 surpassed cases of O157 in Ireland, France, Italy, and Denmark [15]. A serogroup O26 strain, with the *stx2* and *eae* virulence genes, was implicated in multiple HUS cases in young children in France, Italy, and Romania [16–19] and is attributed to a greater number of HUS cases than O157 in the European Union [15]. STEC serogroup O111 was associated with the first non-O157 STEC outbreak in the United States in 1999 causing multiple HC cases and has since then has accounted for 64% of STEC cases in the United States along with other non-O157 STEC [9–11, 20, 21]. HC cases with STEC O103:H2 and Shiga toxin-negative O145:NM infection, linked to venison consumption in Minnesota, United States, were reported in 2010 [22]. Subsequently, major multistate outbreaks with STEC O145 occurred in 2010 and 2012 in the United States linked to contaminated lettuce and ground beef, respectively; the outbreaks resulted in significant HC to HUS cases and hospitalizations [23, 24]. More recently, an STEC O145 outbreak was reported in the United Kingdom in May-June 2024 causing HC to HUS in infected individuals [25].

Despite the increased reporting of non-O157 STEC infections in recent years, there is limited research on the dynamics of non-O157 STEC colonization of the bovine reservoir especially adaptation to the first bovine gastrointestinal compartment encountered, the rumen. In the context of the rumen, currently available literature is based primarily on *in vitro* studies in rumen fluid. For instance, one *in vitro* study in 2012 found variations in the survival of O157 and non-O157 STEC in bovine rumen fluid from steers on total mixed rations [26]. Compared to O157, serotype O111:H8 displayed a significant increase in growth in rumen fluid after 1 and 2h, serotypes O145:H28 and O103.:H8 had a decrease in growth after 8 and 24h, and serotypes O26:H11 and O111:H8 had similar survival rates as O157, but less than that of the other serotypes, suggesting variations in growth are in response to environmental stressors related to colonization of the bovine gastrointestinal tract [26]. A second *in vitro* study examined the effect of diet on survival of STEC in rumen fluid from sheep and found that non-O157 STEC serotypes (O91:H10, OX3:H-), of both human and bovine origin, persist in the rumen fluid in a similar manner despite low oxygen and nutrient availability [27]. The rumen fluid collected from animals on a hay diet contained higher acetate concentrations than rumen fluid collected from animals on a hay and corn diet. Both the human and bovine STEC isolates were shown to utilize acetate as the carbon source to grow in rumen fluid under anaerobiosis [27].

Previously, we developed a non-terminal, reusable, live-animal model to study microbes *in vivo* in the rumen that utilized rumen-fistulated cattle with specialized cartridges to expose bacteria to the rumen environment in a manner that did not inoculate the animal. We used this model to determine growth and protein expression characteristics of different O157 strains [28]. Our comparative research indicated that O157 expresses proteins involved primarily in survival rather than virulence in rumen fluid from animals on different diets, both *in vitro* and *in vivo* [28, 29]. The O157 strains also exhibited distinct *in vitro* but shared *in vivo* growth patterns that grouped them apart from the control non-STEC *E. coli* Nal^R^ [28]. To expand on these inferences, we utilized the same animal model and similar growth conditions to evaluate non-O157 STEC survival and protein expression. We conducted a comparative study investigating growth and protein expression of three non-O157 serotypes O26:H11, O111:H8 and O145:NM along with the control non-STEC *E. coli*: (i) *in vitro* in rumen fluid from cattle on the maintenance (high fiber, low protein) or the lactation (low fiber, high protein) diets, at 39°C for 48 h under anaerobic conditions, and (ii) *in vivo* for 48 h within the fistulated rumen of cattle on the similar diets using the non-terminal reusable-animal model.

## MATERIALS AND METHODS

### Bacterial strains

Four different *E. coli* strain/serotypes with diverse combinations of the Shiga toxin and intimin encoding genes were used in this study including, **(i)** *E. coli* Nal^R^ (*stx1*-, *stx2*-, *eae*-), a non-pathogenic, nalidixic acid-resistant derivative of a bovine commensal *E. coli* (National Animal Disease Center (NADC), Ames, IA), **(ii)** Serotype O26:H11 (DEC10B, *stx_1_* +, *stx_2_*-, *eae*+), a patient isolate associated with HC (STEC Center, Michigan State University, East Lansing, MI), **(iii)** Serotype O111:H8 (DEC8B, *stx_1_* +, *stx_2_*+, *eae*+), a patient isolate associated with HC (STEC Center), and **(iv)** Serotype O145:NM (*stx_1_* -, *stx_2_*-, *eae*-), a porcine diarrheal isolate in the NADC collection.

### Rumen fistulated cattle and diet

#### (i) Animals

Protocols approved by the Institutional Animal Care and Use Committee at the NADC were used. A total of four rumen-fistulated Holstein heifers/cows (#A - #D), 3 - 4 years of age and routinely used as rumen fluid donors at the NADC, were utilized for this project. All experiments were conducted in duplicate. For the *in vitro* experiments, rumen fluid was collected from animals #A and #D, on the lactation and maintenance diets respectively, to evaluate the four different *E. coli* strains. For the *in vivo* experiments, animals #A and #B were used and bacteria exposed to the rumens of both animals, sequentially, to factor in effects of any *in vivo* host-related variations on bacteria being tested.

Both animals were on the lactation diet for the first half of the *in vivo* experiments, then transitioned and acclimated to the maintenance diet over 2 weeks to keep them on this maintenance diet for the second half. Animal #C, used as a control for the *in vivo* experiments, remained unexposed to the test bacteria but experienced all the dietary changes along with the test animals to comparatively monitor diet-related changes in rumen fluid. Feed intake and body temperatures were monitored daily for all animals. Animals were housed in barns at the NADC; in a field barn initially and subsequently in BSL-2 containment barns when ready to introduce the *E. coli* strains into the rumen. At the end of the study, following confirmation of their negative-shedding status for all *E. coli* strains tested, all animals were returned to the field barn.

#### (ii) Diet

As in the previous study [28] the animals were fed one of two diets typically provided to dairy cattle: the lactation diet (L diet) usually provided at the time of lactation to meet the high energy needs of the cow, or the total mixed ration maintenance diet (M diet) usually formulated to maintain the overall weight of the animal when dry [30, 31]. All dietary compositions were monitored by an animal nutritionist on site and prepared as reported previously [28]. The M diet comprising 25% grass hay, 65% corn silage, and 10% Steakmaker 40-20 (Fig. S1; Land O’Lakes, Inc., Arden Hills, MN) was limit fed at ∼19 lbs/head with access to pasture or at ∼30 lbs/head without pasture. The L diet with 2.5% grounded corn, 62% corn silage, 2.5% soybean meal, 12% legume hay and 21% lactation premix (Fig. S1) was limit fed at ∼99 lbs/head. The cattle were initially fed the L diet for 2 weeks following which, when needed, the diet was gradually switched to the M diet over 2 weeks to allow for acclimation. The M diet was then continued for another 2 weeks. Water was provided ad libitum throughout.

### Bacterial culture

#### (i) Inoculum preparation

Log-phase cultures of each bacterial strain were prepared in Luria-Bertani (LB) broth, with or without nalidixic acid (100 μg/ml; LB-Nal), at 39°C with aeration, as described previously [28]. Bacteria harvested from the log-phase cultures at an OD_600_ 0.5-0.6, washed and re-suspended in sterile phosphate-buffered saline (PBS), were used to charge dialysis cartridges as described below under “*Evaluation of STEC strains*”. All STEC cultures were confirmed by culture on Rainbow Agar O157 (Biolog, Hayward, CA) as described below, and serologically by using latex agglutination tests (*E. coli* O157 latex, Oxoid Diagnostic Reagents, Oxoid Ltd., Hampshire, UK and *E. coli* non-O157 identification kit, Pro-Lab Diagnostics, Ontario, Canada).

#### (ii) Isolation of bacteria from dialysis cartridges

Post-incubation, bacteria adhering to the dialysis cartridge walls were scrapped off, using sterile inoculum loops, into the suspension within before aspirating the same. Serial dilutions of the aspirated bacterial suspensions were prepared using sterile 0.9% saline and plated on LB-Nal and/or sorbitol MacConkey (BD Biosciences) with 4-methylumbelliferyl-β-d-glucuronide (MUG, 100 mg/liter; Sigma; SMAC-MUG) agar to determine viable colony counts in CFU/ml. Well-isolated sorbitol- fermenting, and MUG-utilizing (fluorescent under UV light; O26 and O145) or non-utilizing (non-fluorescent; O111) colonies on SMAC-MUG were individually plated on Rainbow Agar O157 (Biolog). The Rainbow Agar O157 plates were incubated overnight at 37°C and colonies were selected based on color for further serological verification of the serogroup as follows: O157- black; O26 – purple/magenta; O111 – bluish-gray; O145 – purplish-gray and *E. coli* Nal^R^- purplish-pink colonies (Biolog; [32, 33]).

#### (iii) Rumen fluid and fecal sample testing for O157 and non-O157 STEC

As done previously, rumen fluid and fecal samples were cultured over two weeks, to determine the presence of any STEC in the animal, both at the beginning and end of the study [28]. Standardized non-enrichment and selective-enrichment culture protocols were used with modifications [34–36]. Briefly,10 ml rumen fluid or 10-g fecal sample was added to 50 ml Trypticase soy broth (BD Bioscience, San Jose, CA) supplemented with cefixime (50 μg/liter; U.S. Pharmacopeia, Washington D.C), potassium tellurite (2.5 mg/liter; Sigma-Aldrich Corp., St. Louis, MO), and vancomycin (40 mg/liter; Alfa Aesar, Haverhill, MA) (TSB-CTV) or LB-Nal broth and mixed well. Serial dilutions of each sample were prepared with sterile 0.9% saline both before and after overnight incubation of the TSB-CTV or LB-Nal suspension at 37°C with aeration. The dilutions prepared before incubation (non-enrichment cultures) and after overnight incubation (selective-enrichment cultures) were spread plated onto SMAC-MUG and LB-Nal plates. SMAC-MUG plates were read after overnight incubation at 37°C and colonies that did not ferment sorbitol or utilize MUG (non-fluorescent under UV light) were further evaluated to be O157 serologically. Fluorescent (MUG-utilizing) and non-fluorescent colonies that fermented sorbitol were cultured on the Rainbow Agar O157 plates and serologically tested for O26, O111 and O145 serogroups as described above.

### Rumen fluid preparation and analysis

Rumen fluid samples collected from rumen-fistulated cows, fed the L diet (LRF) or the M diet (MRF), were used for culture and to set up the *in vitro* experiments for this study. As described previously, up to five liters of either rumen fluid, collected 2–3 hr post-feeding to allow for rumination to occur, was strained through cheesecloth to remove large feed particles and poured into collection flasks [28, 29].

Following recording of the pH and freezing of aliquots for volatile fatty acid (VFA) analysis, the remaining strained rumen fluid was aliquoted into flasks for *in vitro* studies as described below. The pH of the rumen fluid was determined using pH paper (pH range 5.0–8.0; Micro Essential Laboratory Inc., Brooklyn, NY) and VFA concentrations determined by capillary gas chromatography, on an Agilent 6890 N gas chromatograph (Agilent Technologies, Inc., Santa Clara, CA) as described previously [28, 37, 38]. Three technical replicates were used per rumen fluid sample to determine VFA concentrations and of the entire panel of substrates evaluated, concentrations of three VFAs, acetate, propionate, and butyrate, were specifically studied as before since these influence bovine growth and energy dynamics [28, 30, 31, 39]. The rumen fluid pH and VFA were recorded before and after introduction of STEC strains at the end of each incubation, in both *in vitro* and *in vivo* studies. Intermittent sampling was avoided to prevent repeated exposure of rumen fluid/rumen to the external environment.

### Evaluation of STEC strains

All experiments were done in duplicate. Since the primary goal was to determine the influence of diet on bacterial gene expression in the rumen, steps were taken to minimize any host-related effects by alternating the animals used for the *in vivo* studies between the first and second challenges. On the other hand, the *in vitro* experiments were only for comparative purposes, hence rumen fluid from any one animal was used.

#### (i) *In vitro* evaluation in rumen fluid

Rumen fluid was collected from two different rumen-fistulated animals, one on the L diet (animal #A) and the other on the M diet (animal #D), for the respective *in vitro* experiments, and distributed separately in 300 ml aliquots per sterile flasks with rubber stoppers. To create anaerobic culture conditions, as described previously, the flasks were transferred into the anaerobic Coy Chamber for 72 hr, charged with bacteria containing dialysis cartridges within the chamber, sealed and then removed for incubation at 39°C for 48 hr [28]. Sterile dialysis cartridges (Ultra-test Dialyzer Chamber, Harvard Apparatus, Hollister, MA), charged with the 4 ml of the bacterial suspension in PBS and sealed at both ends with polycarbonate membranes (0.01um pore size, Harvard Apparatus), were used to contain the test bacteria within allowing for easy recovery while permitting diffusion of rumen fluid through the cartridge [28]. A total of 4 dialysis cartridges per bacterial strain were used.

#### (ii) *In vivo* evaluation in the rumen

Two rumen-fistulated animals, #A and #B, were used to set up the *in vivo* studies; both animals were fed the lactation diet in the first half of the *in vivo* study and the maintenance diet in the second half, post-acclimation. In each half of the study, the test bacteria were exposed to both rumens sequentially, by alternating the animal used for the experiment, to account for possible host-related influences on the bacteria besides the diet itself. A total of 4 dialysis cartridges per bacterial strain were used. As done previously, the dialysis cartridges charged with bacteria were placed in mesh laundry bags tethered to a 3 ft. fishing line attached to the rumen cannula cap to ensure retrievability from the rumen [28]. All bags with the charged dialysis cartridges were introduced into the rumen and retrieved only after 48 hr, then transported to the lab on ice where the integrity of the dialysis cartridges was verified and contents processed [28].

### Proteomics

#### (i) Protein sample preparation and labelling for Isobaric Tags for Relative and Absolute Quantification (iTRAQ) analysis

Following each *in vitro* and *in vivo* experiment, harvested bacteria of the same serogroup and growth condition were pelleted together. The pellets were washed three times with ice-cold sterile PBS (pH 7.4), frozen and subsequently processed to obtain bacterial cellular and membrane proteins for proteomic analysis as described previously [28, 40]. For iTRAQ, the protein fractions were quantified [41] and dissolved in denaturant buffer (0.1% SDS (w/v) and dissolution buffer (0.5 M triethylammonium bicarbonate, pH 8.5) in the iTRAQ 8-plex kit (AB SCIEX Inc., Foster City, CA). A total of 100 μg of protein per sample were reduced, alkylated, trypsin-digested, and resulting peptides labeled with either of the iTRAQ tags 114, 115, 116,117, 118, 119 or 121, according to the manufacturer’s instructions (AB SCIEX Inc.). Peptides from bacteria grown in MRF-*in vitro* and MRF-*in vivo* were labeled separately based on strain-growth condition and then mixed in one tube; the LRF-*in vitro* and LRF-*in vivo* peptides were processed likewise. The mixed labeled peptides were dried and held at -80°C until ready for processing as follows. The labeled peptides were desalted with C18-solid phase extraction and dissolved in strong cation exchange (SCX) solvent A (25% (v/v) acetonitrile, 10 mM ammonium formate, and 0.1% (v/v) formic acid, pH 2.8).

#### (ii) Strong cation exchange fractionation and reverse-phase liquid chromatography with tandem mass spectrometry (LC-MS/MS)

The peptides were fractionated using an Agilent HPLC 1260 with a polysulfoethyl A column (2.1 × 100 mm, 5 µm, 300 Å; PolyLC, Columbia, MD). Peptides were eluted with a linear gradient of 0–20% solvent B (25% (v/v) acetonitrile and 500 mM ammonium formate, pH 6.8) over 50 min followed by ramping up to 100% solvent B for 5 min. The absorbance at 280 nm was monitored and a total of 10 fractions per sample were collected. The fractions were lyophilized and resuspended in LC solvent A (0.1% formic acid in 97% water (v/v), 3% acetonitrile (v/v)). A hybrid quadrupole Orbitrap (Q Exactive Plus) MS system (Thermo Fisher Scientific, Bremen, Germany) was used with high energy collision dissociation (HCD) in each MS and MS/MS cycle. The MS system was interfaced with an automated Easy-nLC 1000 system (Thermo Fisher Scientific). Each sample fraction was loaded onto an Acclaim Pepmap 100 pre-column (20 mm × 75 μm; 3 μm-C18) and separated on a PepMap RSLC analytical column (250 mm × 75 μm; 2 μm-C18) at a flow rate of 350 nl/min during a linear gradient from solvent A (0.1% formic acid (v/v)) to 30% solvent B (0.1% formic acid (v/v) and 80.0% acetonitrile (v/v)) for 95 min, to 98% solvent B for 15 min, and hold 98% solvent B for an additional 30 min. Full MS scans were acquired in the Orbitrap mass analyzer over m/z 400–2000 range with resolution 70,000 at 200 m/z. The top ten most intense peaks with charge state ≥ 2 were isolated (with 2 m/z isolation window) and fragmented in the high energy collision cell using a normalized collision energy of 28% selected. The maximum ion injection time for the survey scan and the MS/MS scans were 250 ms, and the ion target values were set to 3e6 and 1e6, respectively. The selected sequenced ions were dynamically excluded for 60 sec.

### Analysis of the iTRAQ data

#### (i) Database searching

Tandem mass spectra were extracted by Proteome Discoverer (Thermo-Fisher) version 2.2.0.388. Charge state deconvolution and deisotoping were not performed. All MS/MS samples were analyzed using Mascot (Matrix Science, London, UK; version 2.4.1). Mascot was set up to search the *E. coli* O26:H11, *E. coli* O111:H8, and *E. coli* O145:NM databases, that are also part of the *E. coli* pan proteome, individually, assuming the digestion enzyme trypsin. Mascot was searched with a fragment ion mass tolerance of 0.60 Da and a parent ion tolerance of 20 PPM. Methylthio of cysteine and iTRAQ8plex of lysine and the n-terminus were specified in Mascot as fixed modifications. Gln->pyro-Glu of the n-terminus, deamidated of asparagine and glutamine, methyl of lysine, oxidation of methionine, phospho of serine, threonine and tyrosine, nmethylmaleimide of cysteine and iTRAQ8plex of tyrosine were specified in Mascot as variable modifications.

#### (ii) Criteria for protein identification

Scaffold (version Scaffold_4.11.1, Proteome Software Inc., Portland, OR) was used to validate MS/MS based peptide and protein identifications. Peptide identifications were accepted if they could be established at greater than 80 - 95% probability by the Peptide Prophet algorithm [42] with Scaffold delta-mass correction. Protein identifications were accepted if they could be established at greater than 80 - 95% probability and contained at least 2 identified peptides. False discovery rate (FDR) was at medium confidence; 0.2 - 0.5% for proteins post-MR exposure and 2.5 - 3% for proteins post- LRF exposure. Protein probabilities were assigned by the Protein Prophet algorithm [43].

Proteins that contained similar peptides and could not be differentiated based on MS/MS analysis alone were grouped to satisfy the principles of parsimony. Proteins sharing significant peptide evidence were grouped into clusters.

#### (iii) Intensity Based Absolute Quantification (iBAQ) analysis of the LC- MS/MS data

MaxQuant [44] was used for label-free peptide and protein identification, protein iBAQ quantification, and reporter ion intensity quantification. The Uniprot *E. coli* pan proteome UP000000625 [45] was used as the search database. Only proteins identified by more than one peptide with Q values less than 0.05 were considered for these analyses. iBAQ values comparing the abundance of proteins between the two diets were transformed into relative iBAQ (riBAQ) values (Fig. S2). Each protein was represented by a single riBAQ value within each diet. To detect proteins which were differentially abundant within each diet the log2FoldChange (L2FC) was calculated by: log2(lact_riBAQ / maint_riBAQ). Proteins with an absolute L2FC of greater than 1 were considered differentially abundant.

#### (iv) iTRAQ quantitation without reference comparison

Two different methods were tested to normalize reporter ion intensities. NOMAD, normalization of mass spectrometry data, was used to normalize peptide reporter ion intensities and aggregate peptide reporter ion intensities to protein level intensities. This is an ANOVA type of normalization and is similar to what can be performed in Scaffold. In addition, centered log ratios (clr) were used to normalize aggregated protein intensities. This is a more general normalization that can be performed on many different ‘-omics’ data [46]. Differentially abundant proteins between *in vivo* and *in vitro* conditions were determined using T-tests and P values were corrected by the FDR method.

### Statistics and bioinformatics

The 1-way or 2-way ANOVA and T tests were used to evaluate the VFA concentrations and viable counts, separately, and a value of *p*<0.05 was considered as statistically significant (GraphPad Prism v9; GraphPad Software, Inc., San Diego, CA). The predicted cellular location of all proteins was determined using PSORTb v3.0.2 [40, 47]. Gene Ontology (GO) terms to all proteins were annotated in the reference databases with InterProScan5 [48]. For sets of differentially expressed proteins enriched GO-terms were calculated with the R package TopGO [49–52]. To investigate the multivariate similarity between samples, ‘robust Aitchison’ distances were used. These distances were used to generate NMDS ordinations to visualize the multivariate similarities between samples. For each STEC strain tested, ratios for each protein expressed in LRF or MRF, under *in vivo* and *in vitro* growth conditions, were determined as follows. Using the log_2_(vivo/vitro) calculation, proteins were assigned to greater expression *in vivo* (positive ratio) or *in vitro* (negative ratio) per strain and the data for the top fifty proteins with the most variation between strains were used to generate comparative heatmaps. The parameters used for all MaxQuant-iBAQ, and reference-free iTRAQ data analysis are available (Supplemental File1_iBAQ-code, Supplemental File2_ReferenceFreeiTRAQ-code).

## RESULTS

### The ruminal pH and VFA concentrations in LRF and MRF were primarily influenced by diet

As observed previously, the two diets had differing influences on the ruminal pH and volatile fatty acid (VFA) profiles (Fig. 1: Tables 1A, 1B and Fig. 2: Tables 2A, 2B).

**Figure 1:**
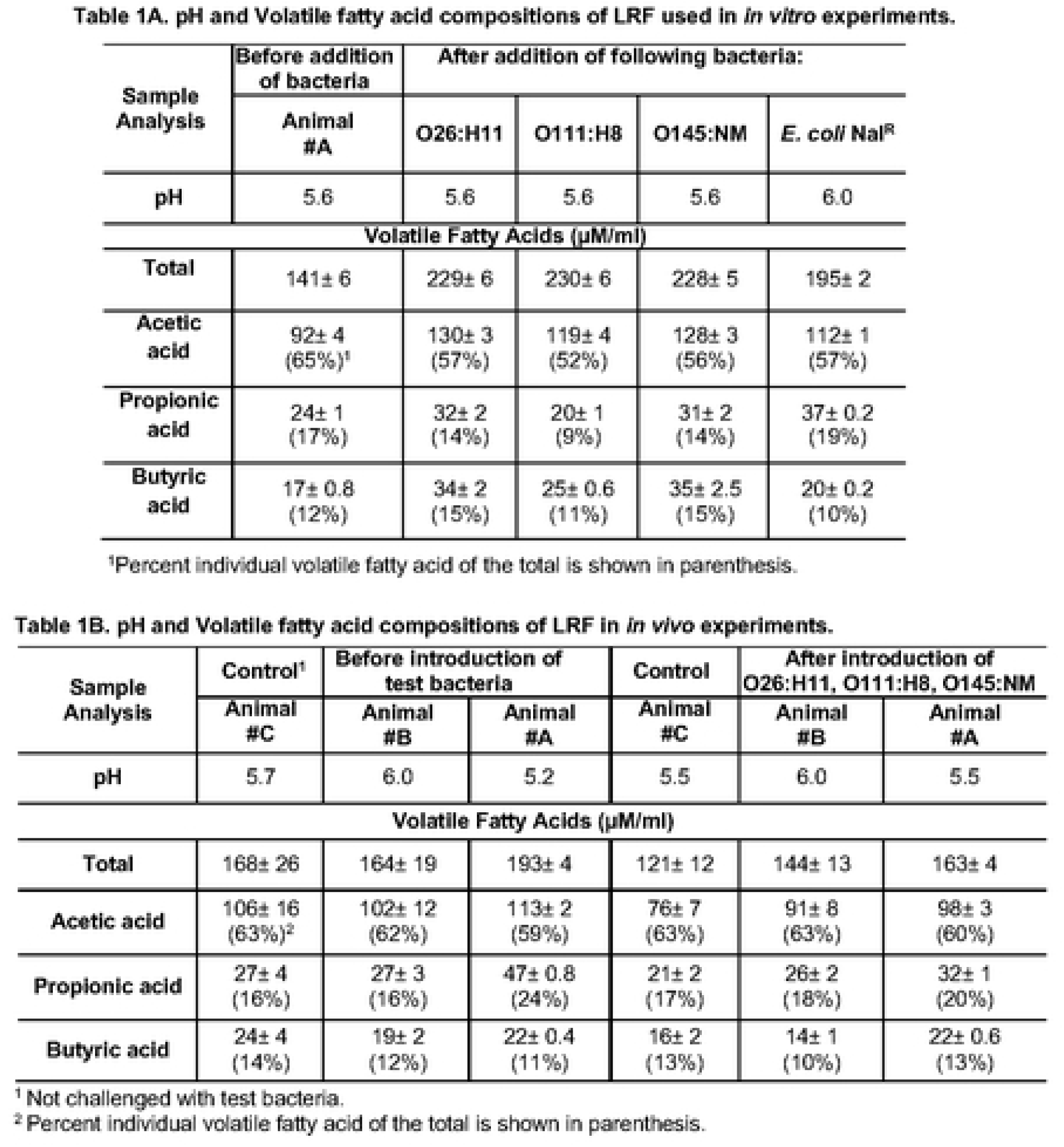
pH and volatile fatty acid compositions of LRF from the *in vitro* (Table 1A) and *in vivo* (Table 1B) experiments.

**Figure 2.**
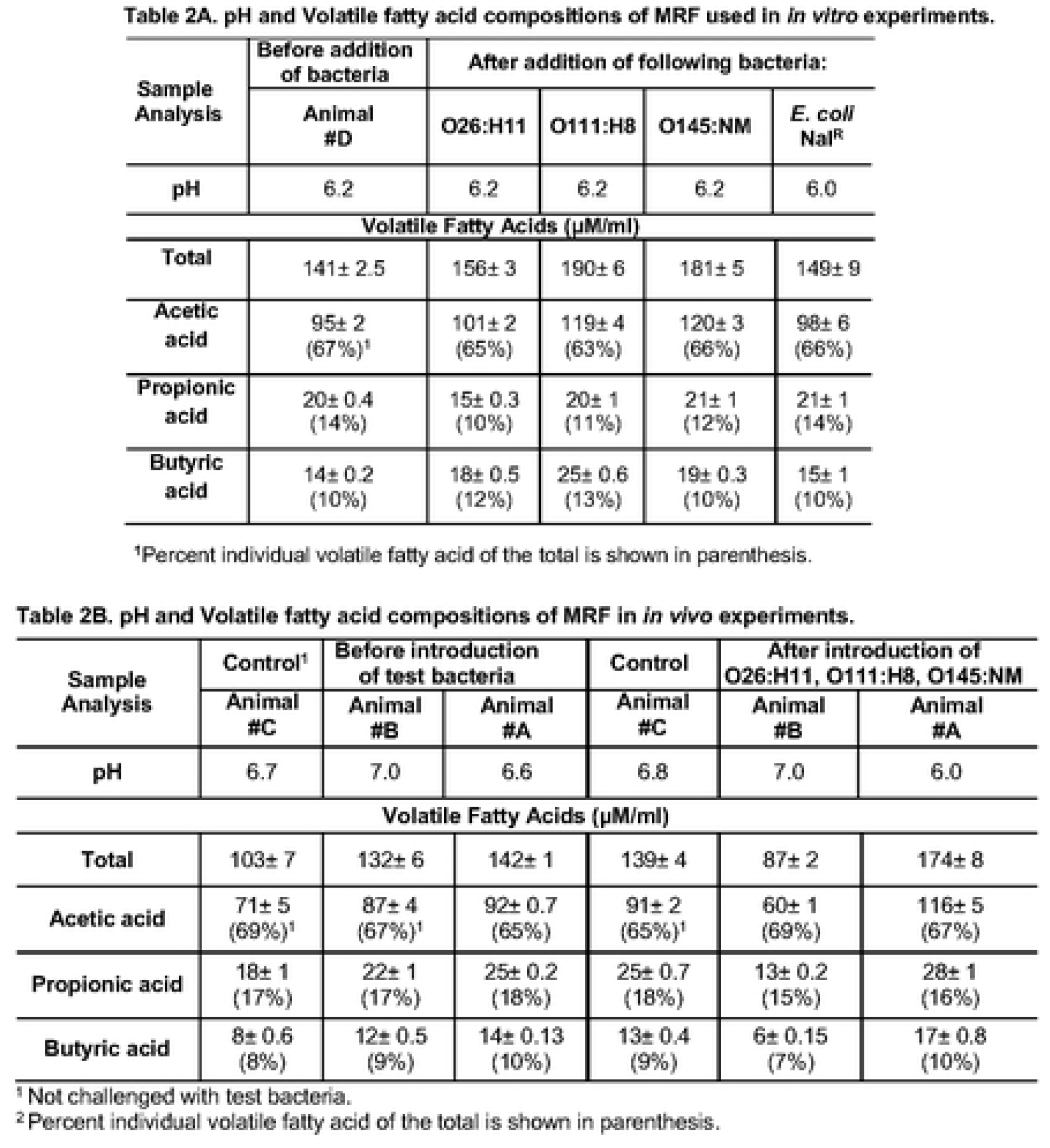
pH and volatile fatty acid compositions of MRF from the *in vitro* (Table 2A) and *in vivo* (Table 2B) experiments.

#### (i) In LRF

*In vitro*, the pH of the rumen fluid collected from animal #A on the L diet was at 5.6 prior to addition of dialysis cartridges with STEC. The rumen fluid pH ranged between 5.6 – 6.0, post-incubation, in the different flasks with STEC containing dialysis cartridges suggesting absence of any bacterial effects (Fig.1: Table 1A). Likewise, no bacterial influence on ruminal pH was observed in the *in vivo* experiments with animals #A and #B on the L diet; pH ranged from 5.2 - 6.0 prior to introduction of dialysis cartridges into the rumen and remained between 5.5 - 6.0 after 48 h (Fig.1: Table 1B). The ruminal pH of the control animal #C ranged between 5.7 and 5.5 at timepoints matching pre- and post- challenge of the test animals (Fig. 1: Table 1B).

The total VFA concentrations ranged from 141 - 230 μM/ml in the *in vitro* samples with higher values occurring post-exposure to test bacteria (Fig. 1: Table 1A). *In vivo*, the ruminal total VFA concentrations were between 144 - 193 μM/ml for the test animals and between 121 - 168 μM/ml for the control animal (Fig. 1: Table 1B). In contrast to the observed increase in VFA post-bacterial exposure *in vitro*, VFA values were lower *in vivo* post-exposure to the test bacteria. Since this decrease was also observed with the control animal it is unlikely this was a bacterial effect, and more host related.

### (ii) In MRF

In the *in vitro* experiments, the pH of the rumen fluid collected from animal #D on the maintenance diet ranged from 6.0 to 6.2 irrespective of the presence of the test bacteria (Fig. 2: Table 2A). The total VFA concentrations ranged between 141 - 190 μM/ml (Fig.2: Table 2A), with increased concentrations observed post-exposure to test bacteria as with LRF-*in vitro*. In the *in vivo* experiments, the ruminal pH ranged between 6.0 – 7.0 and total VFA between 87 - 174 μM/ml (Fig2: Table 2B). The test bacteria did not impact the pH or VFA concentrations in the *in vivo* experiments with similar changes observed in the control animal (Fig. 2: Table 2B).

Overall, butyrate concentrations were higher in LRF than MRF while the opposite was true for acetate concentrations, as expected [28, 30, 31]. The differences in the total VFA concentrations, between the pre- and post-exposure samples, were significant (*p*<0.05) in all instances except for the LRF-*in vivo* samples.

### Serogroup O26, O111 and O145 strains exhibited similar survival patterns in LRF and MRF

Contrary to the observations with the STEC O157 strains [28], the non-O157 serotypes tested had similar survival patterns, both *in vitro* and *in vivo,* that were distinct from the non-STEC *E. coli* Nal^R^ only *in vitro* (Figs. 3 and 4).

**Figure 3.**
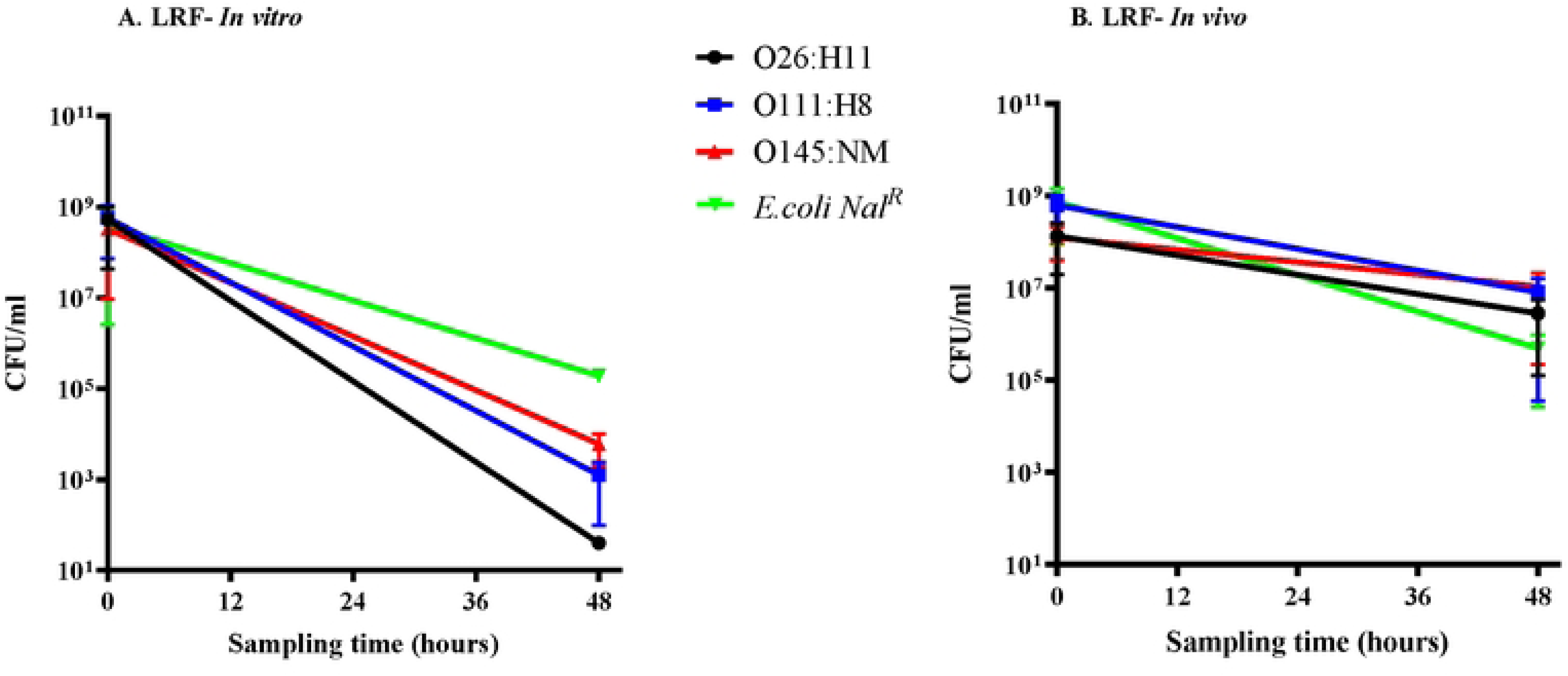
Graphs shown represent survival characteristics of the three serotypes in comparison to the control *E. coli* Nal^R^, in (A) *in vitro* and (B) *in vivo* assays, in LRF. Bacterial survival characteristics depicted are following anaerobic incubation for 48 h, *in vitro* in flasks with LRF or *in vivo* in the rumen of animals fed the lactation diet. Viable counts in colony forming units [CFU]/ml, with the standard error of means, against the sampling time in hours are shown in both graphs. Key for each STEC serotype tested is also shown.

**Figure 4.**
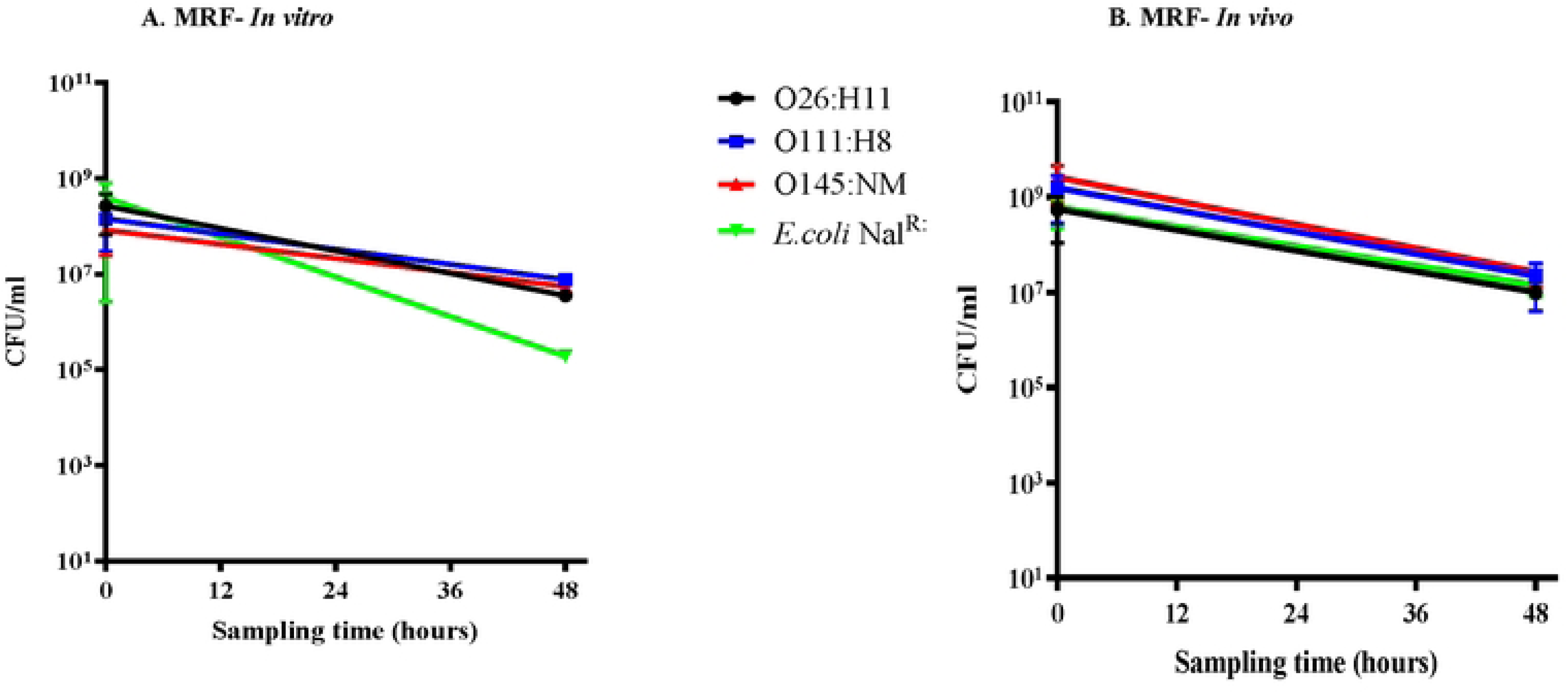
Graphs shown represent survival characteristics of the three serotypes in comparison to the control *E. coli* Nal^R^, in (A) *in vitro* and (B) *in vivo* assays, in MRF. Bacterial survival characteristics depicted are following anaerobic incubation for 48 h, *in vitro* in flasks with MRF or *in vivo* in the rumen of animals fed the maintenance diet. Viable counts in colony forming units [CFU]/ml, with the standard error of means, against the sampling time in hours are shown in both graphs. Key for each STEC serotype tested is also shown.

#### (i) *In vitro* survival in rumen fluid

Post-incubation in LRF, the STEC serogroups demonstrated greater reduction (5 to 7-log) in average viable counts compared to *E. coli* Nal^R^ (3-log), with STEC O26:H11 having the highest reduction by 7-log (Fig. 3; Table S1). In MRF, the STEC serogroups had 2-log reduction in average viable counts post-incubation compared to the 3-log reduction in *E. coli* Nal^R^ viable counts (Fig. 4A; Table S2). Based on these results, it appeared that the non-O157 STEC serogroups have different survival dynamics than *E. coli* Nal^R^, *in vitro*, in both rumen fluids (Figs. 3A and 4A) and were more sensitive to the low pH, higher VFA conditions in LRF-*in vitro*. The test bacteria were not recovered from either rumen fluids, pre- or post- exposure to the dialysis cartridges, thereby verifying the containment of the test bacteria within.

#### (ii) *In vivo* survival in the rumen

In contrast to the *in vitro* results, all three STEC serogroups had a survival pattern similar to that of *E. coli* Nal^R^. Post-48 hr incubation within the rumen of fistulated cows fed the L diet, an average of 1 to 3-log reduction in the viable counts was observed among the tested bacteria (Fig. 3B; Table S1). Likewise, a 1 to 2-log reduction in the averaged viable counts was recorded post-48 hr exposure to the rumen of cows on the M diet (Fig. 4B, Table S2). These results further confirm the importance of conducting *in vivo* experiments as previously reported [28]. The test bacteria were not recovered from the rumen fluid and fecal samples of the animals, including the control animal, prior to and post-exposure to the dialysis cartridges containing the test bacteria.

### Animal diet and growth conditions influenced proteins expressed by non-O157 serotypes in LRF and MRF

#### (i) iBAQ analysis highlighted overall differences in proteins expressed in LRF and MRF, irrespective of growth condition

Using iBAQ analysis proteins expressed collectively by STEC O26:H11, O111:H8 and O145:NM, uniquely or differentially, in MRF and LRF were identified (Tables S3 - S6). Since all samples pooled in each MS run belonged to the same rumen fluid type, in both the *in vitro* and *in vivo* culture conditions, iBAQ data for a particular rumen fluid type could be used to estimate the overall differences in protein expression between MRF and LRF. As done previously, to avoid any false interpretations due to differences in intensities, iBAQ values derived from the LC-MS/MS data were converted to riBAQ values to look at relative protein abundance for each rumen fluid type [28] (Fig. S2) without focusing on individual strains or the *in vitro*/*in vivo* growth conditions. A total of 446 bacterial proteins were identified by riBAQ of which 371 proteins were detected in both rumen fluids, 34 were unique to LRF and 41 were unique to MRF (Tables S3 - S6). In addition, any protein with an absolute L2FC value of 1 or greater was considered to be enriched in the respective rumen fluid (Fig. S3); 91 proteins were found to be enriched in LRF (Tables S3 and S5) and 185 enriched in MRF (Tables S3 and S4). These differentially expressed proteins were used to identify GO terms that were significantly enriched in each rumen fluid (Tables S5 and S6).

#### (ii) Reference-free iTRAQ analysis identified proteins enriched by growth conditions in each rumen fluid

Using reference-free iTRAQ data analysis, along with centered log ratios (clr) normalization, a total of 405 and 412 proteins were identified as being differentially expressed by the three serotypes in LRF and MRF, respectively (Fig. 5). In LRF, of the 405 proteins, 25 were more highly expressed in the *in vivo* condition and 37 were more highly expressed in the *in vitro* condition (Fig. 5, Tables S7 and S8). In MRF, 26 of the 412 proteins were more highly expressed in the *in vivo* condition and 41 were more highly expressed *in vitro* (Fig. 5; Tables S9 and S10). NOMAD normalization yielded minimal data except with proteins expressed *in vitro* in LRF and hence, not used in the final analysis (Tables S11-S14). Multivariate similarity analysis aligned the serotype-proteomes more with growth conditions and showed minimal strain-based variations supporting the observed *in vivo* and *in vitro* survival patterns (Figs. S4, 3 and 4). All the same analysis of the top fifty proteins expressed with most variation between strains, in LRF or MRF, did reveal subtle serotype-related differences (Fig. S5; Tables S15-S16).

**Figure 5.**
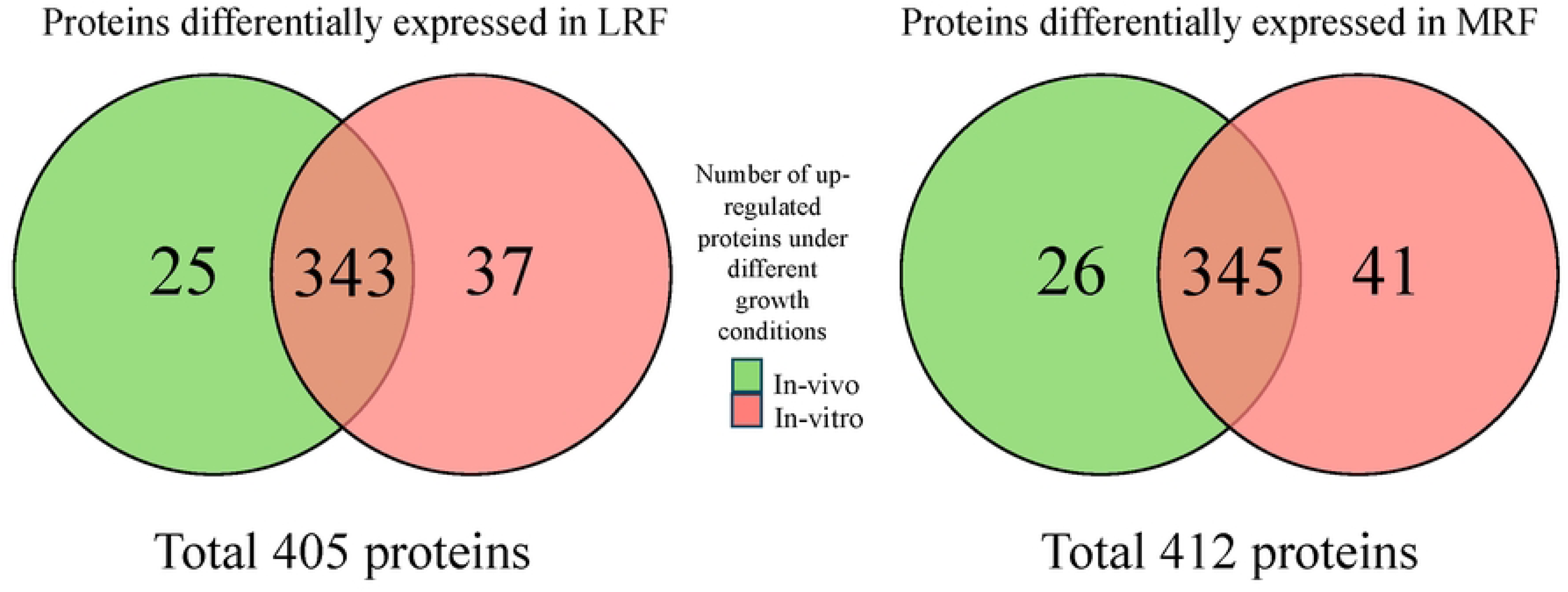
Venn diagrams of total differentially expressed proteins, up regulated *in vivo* and *in vitro*, in LRF and MRF.

#### (iii) iBAQ and iTRAQ analyses provided a broader and in-depth perspective on proteins expressed in LRF and MRF, respectively

Based on the iBAQ analysis, overall, a total of 91 STEC proteins were identified to be enriched in LRF and 185 proteins in MRF (Tables S3 - S6), irrespective of the growth conditions. Analysis of GO-terms for these proteins indicated an enrichment of transport, ribosome assembly, and membrane assembly pathways in LRF and MRF (examples: AcrB, BamD, BtuB, DegP, FadL, FepA, FhuA, HisJ, LamB, OmpF, OmpT, OmpW, SlyB, TolC, WzzB) while biosynthetic pathways were primarily dominant in MRF (Tables S5 - S6). Interestingly, more membrane associated proteins were enriched (L2FC >1) in LRF than in MRF including NmpC, GfcE, LamB, OmpA, CirA, Ag43, Eae, FhuA, FepA, GfcD, BtuB, LptD, OmpX, TolC, BamD, OmpT, BamB, OmpF, SlyB, OmpW, FadL, NuoG, WzzB, PtsG, LpoA, AcrB, PspA, and NarG (Tables S3 - S6). On the other hand, 11 membrane-associated proteins were enriched in MRF, (GdhA, ZipA, SecD, MetQ, FabF2, Alkyl hydroperoxide reductase subunit F, YhcB_1, AtpB, SdhB, FrdA, XdhD) (Tables S3 - S6).

Centered log ratios (clr) normalization helped sort the reference-free-iTRAQ data better than NOMAD (Tables S7 – S14). Following clr normalization, the iTRAQ data could be used to reliably distinguish proteins expressed exclusively *in vivo* or *in vitro*, in both LRF and MRF (Fig. 5, Table 3, S7 – S10). While bacterial proteins identified by iBAQ overlapped with those identified by reference- free-iTRAQ, the data analysis using the latter was nuanced in the context of growth conditions. For instance, membrane-associated proteins expressed in LRF, determined as *in vivo*-expressed included, AtpE, TolC, YdgA, DmsB, BamA, Lpp and those determined as *in vitro*-expressed were FabI, HtpG, AtpF (Tables S7 and S8). Likewise, proteins expressed in MRF were identified as *in vivo*-expressed (FadL, FhuA, LptD, TolC, SlyB) or *in vitro*-expressed (PspA) (Table 3, S9 and S10). All other membrane proteins identified by iBAQ were determined to be expressed under both growth conditions by iTRAQ analysis in either one of the strains tested. Additionally, as observed with iBAQ analysis, proteins associated with metabolic/biosynthetic pathways were enriched *in vitro* and *in vivo* in MRF (Table 3). While virulence-associated proteins were not highlighted by iTRAQ analysis, iBAQ identified proteins that support pathogenicity, namely, intimin (LRF) involved in STEC adherence [53]and glutamine/arginine transport proteins (LRF/MRF) associated with acid resistance [54] (Tables S3-S6).

**Table 3.**
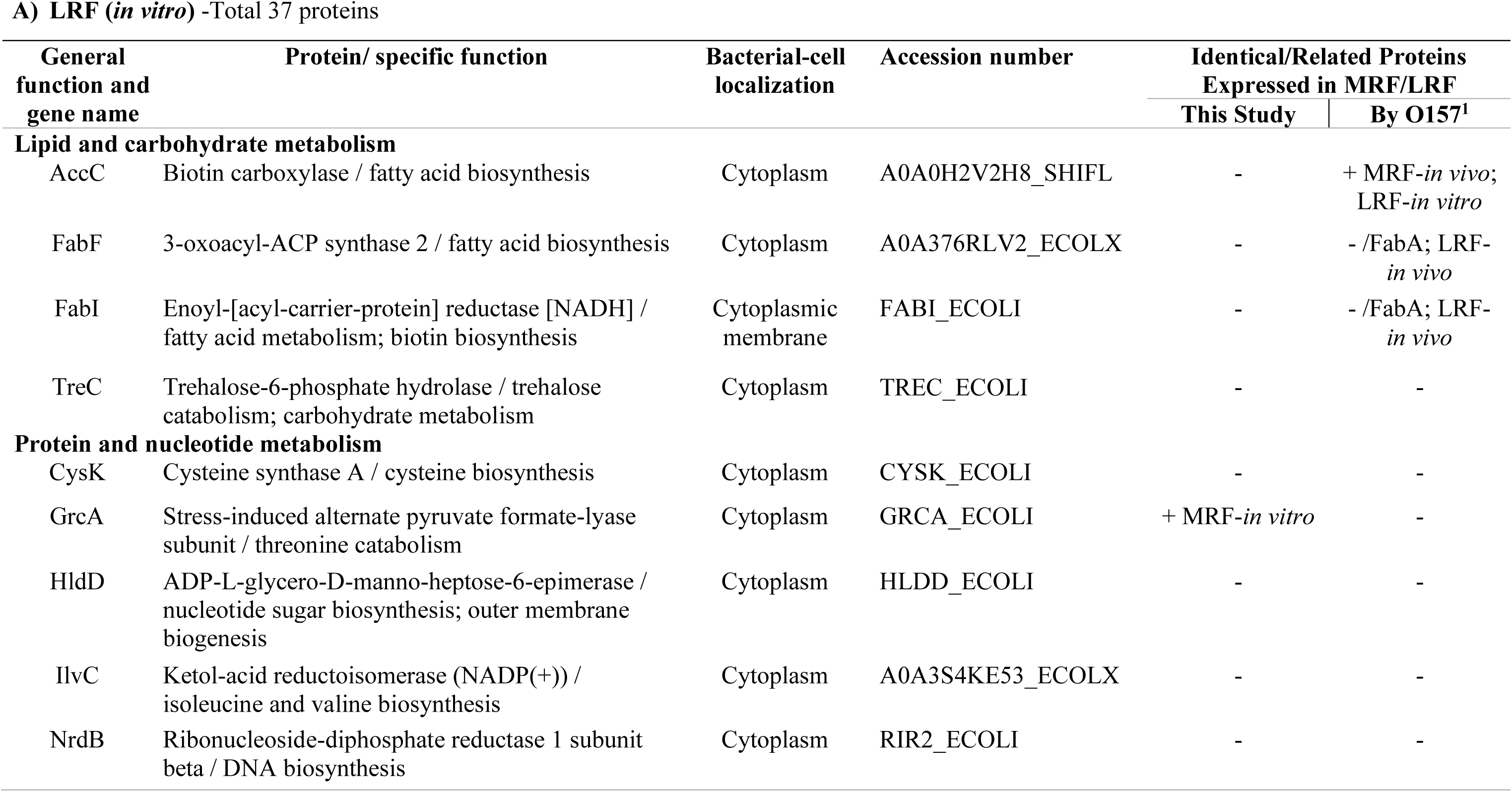
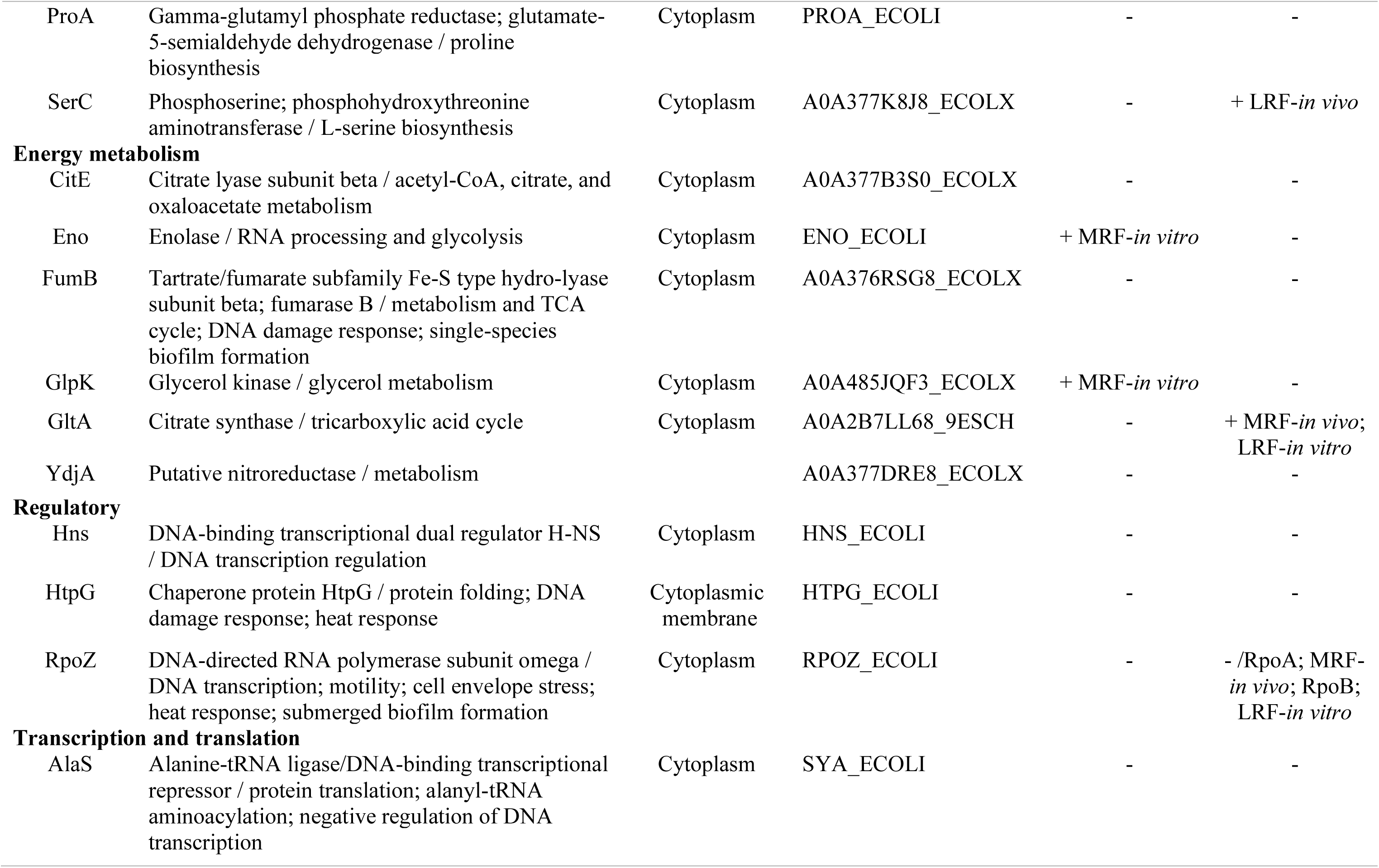
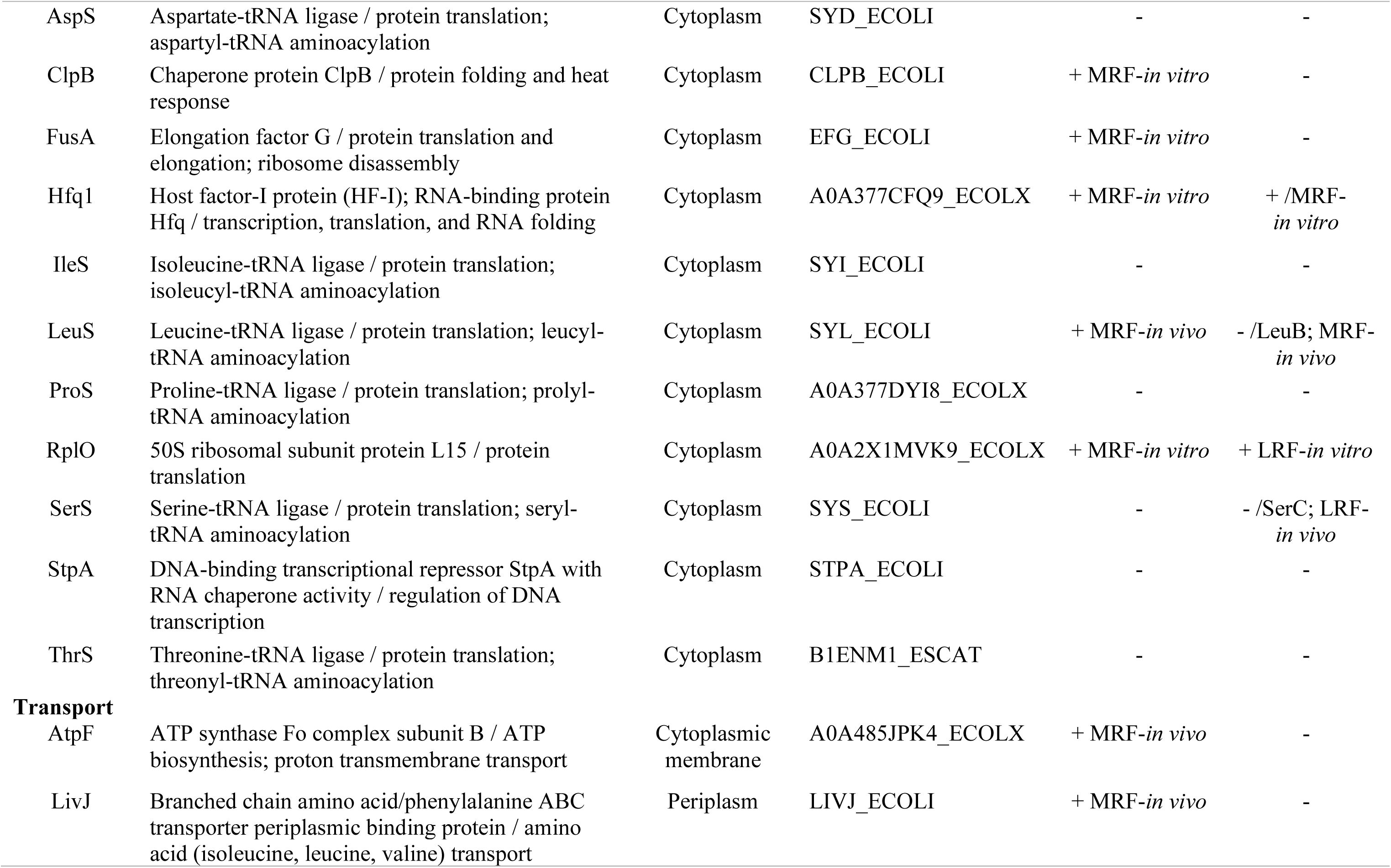
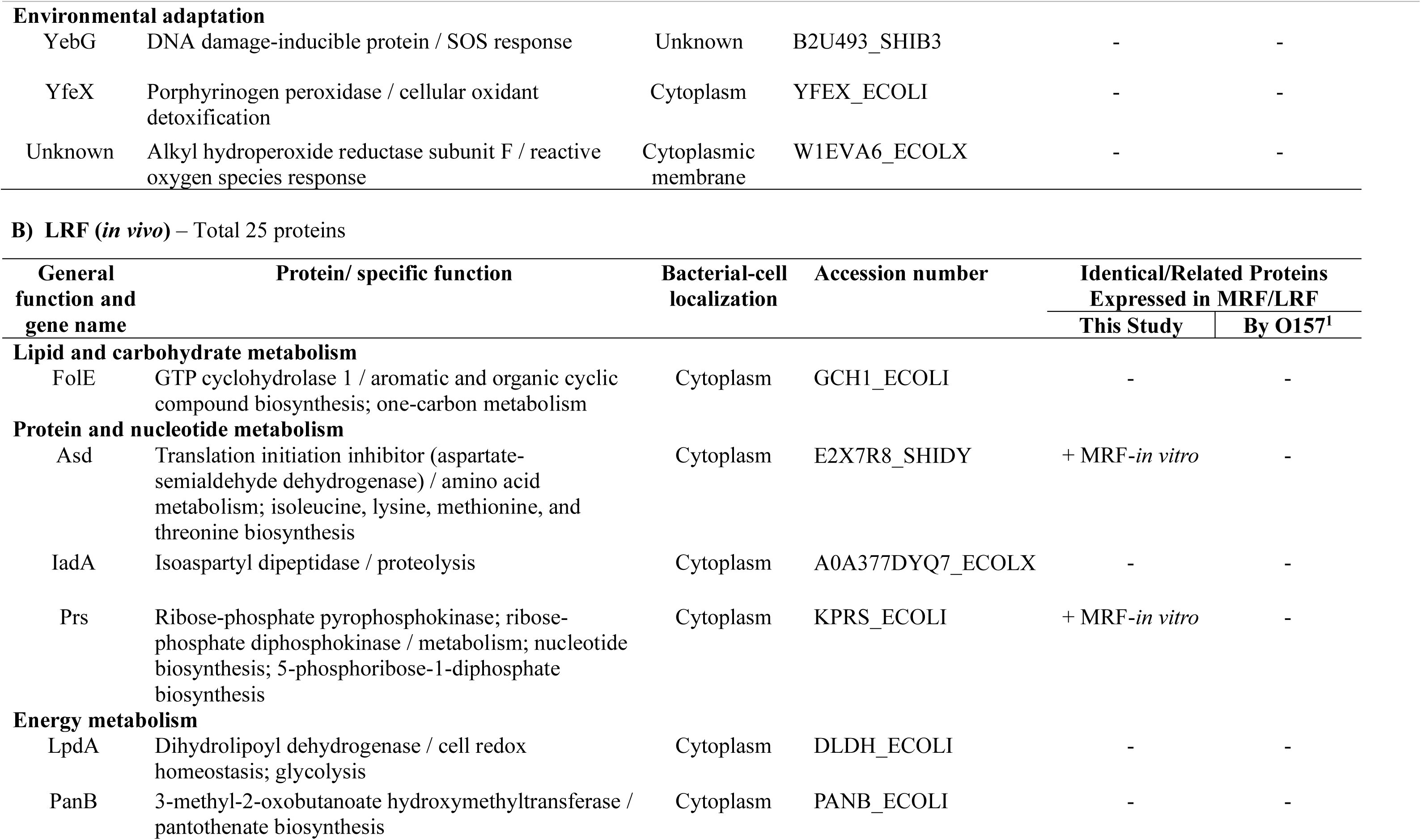
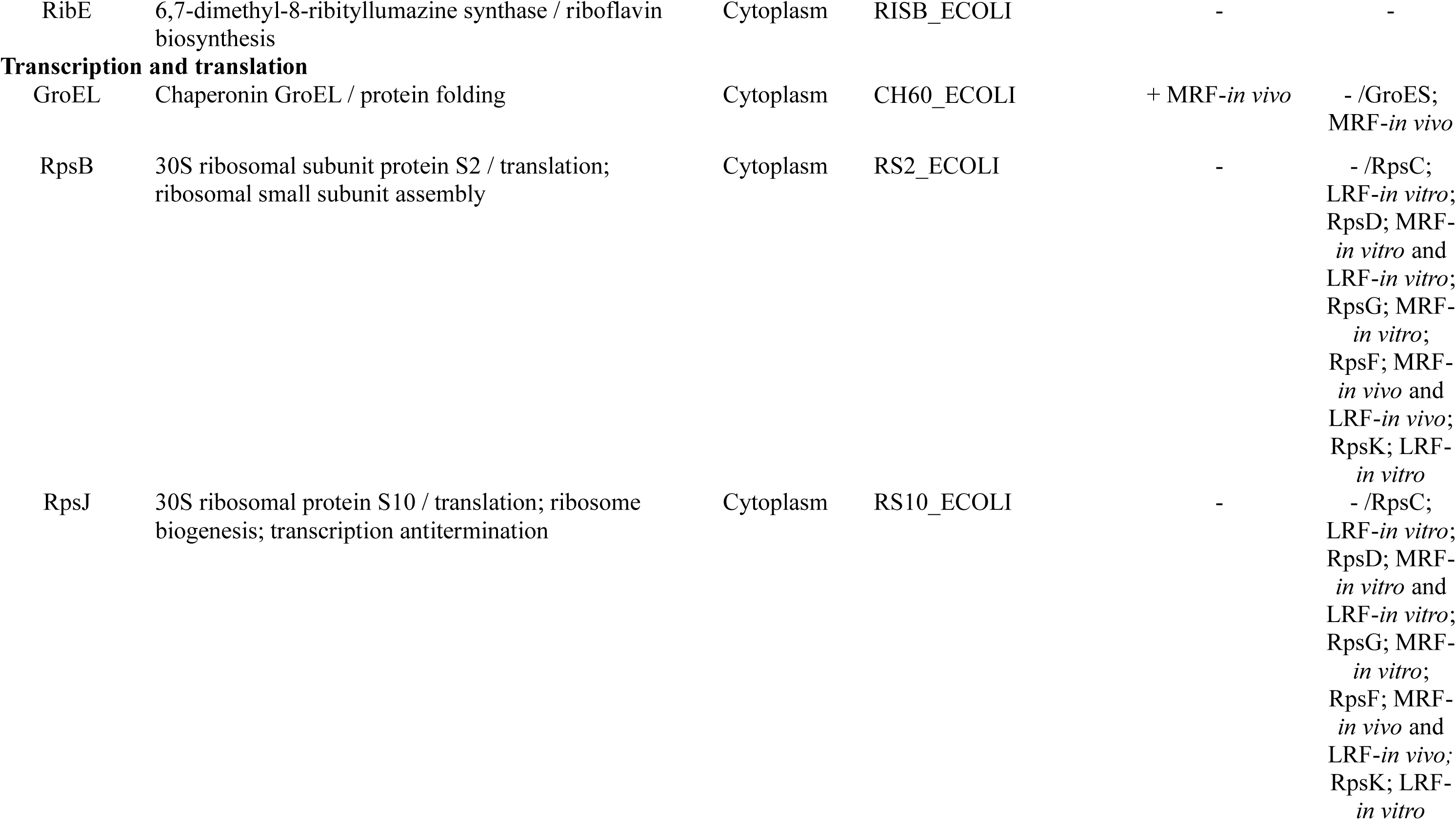
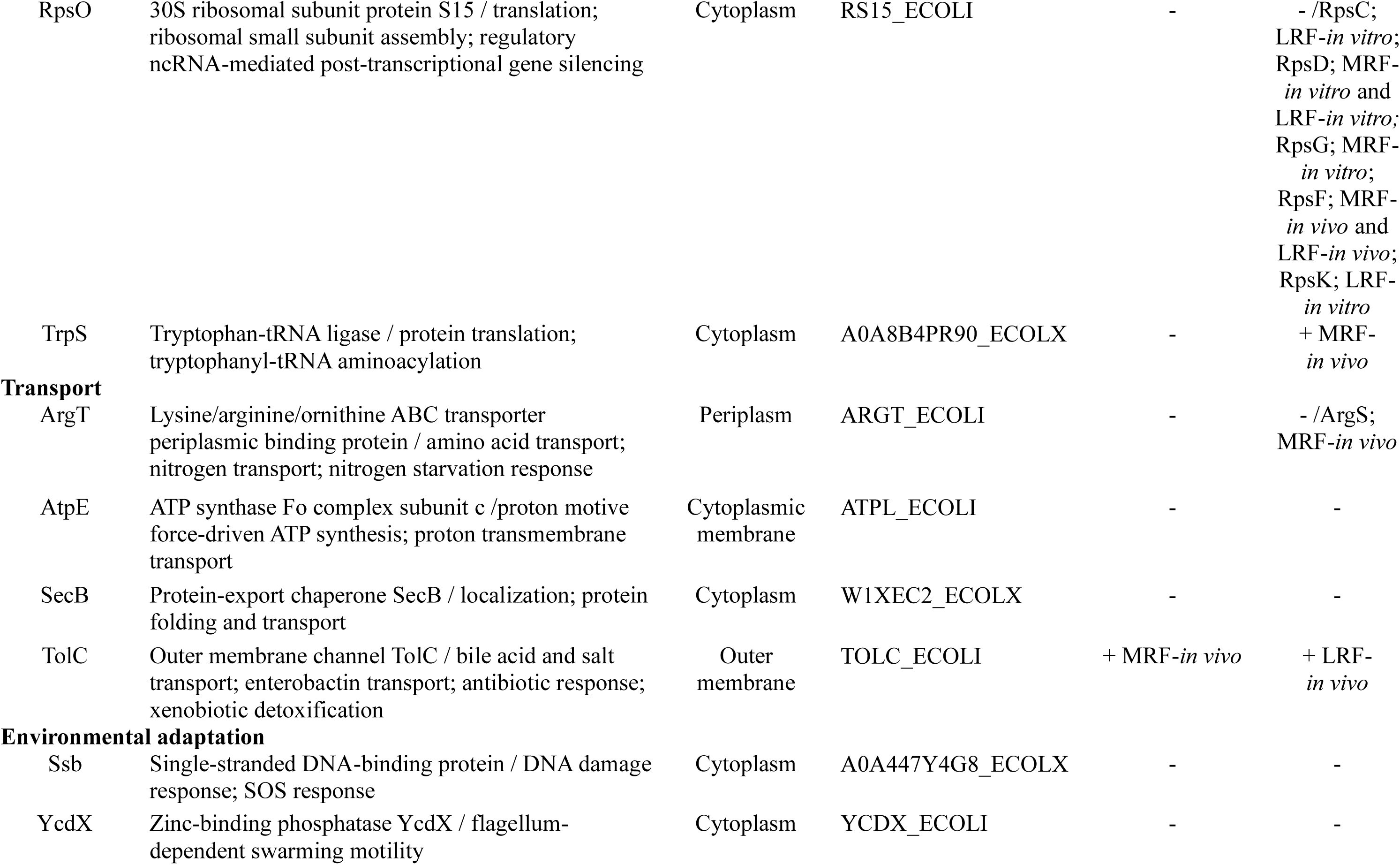
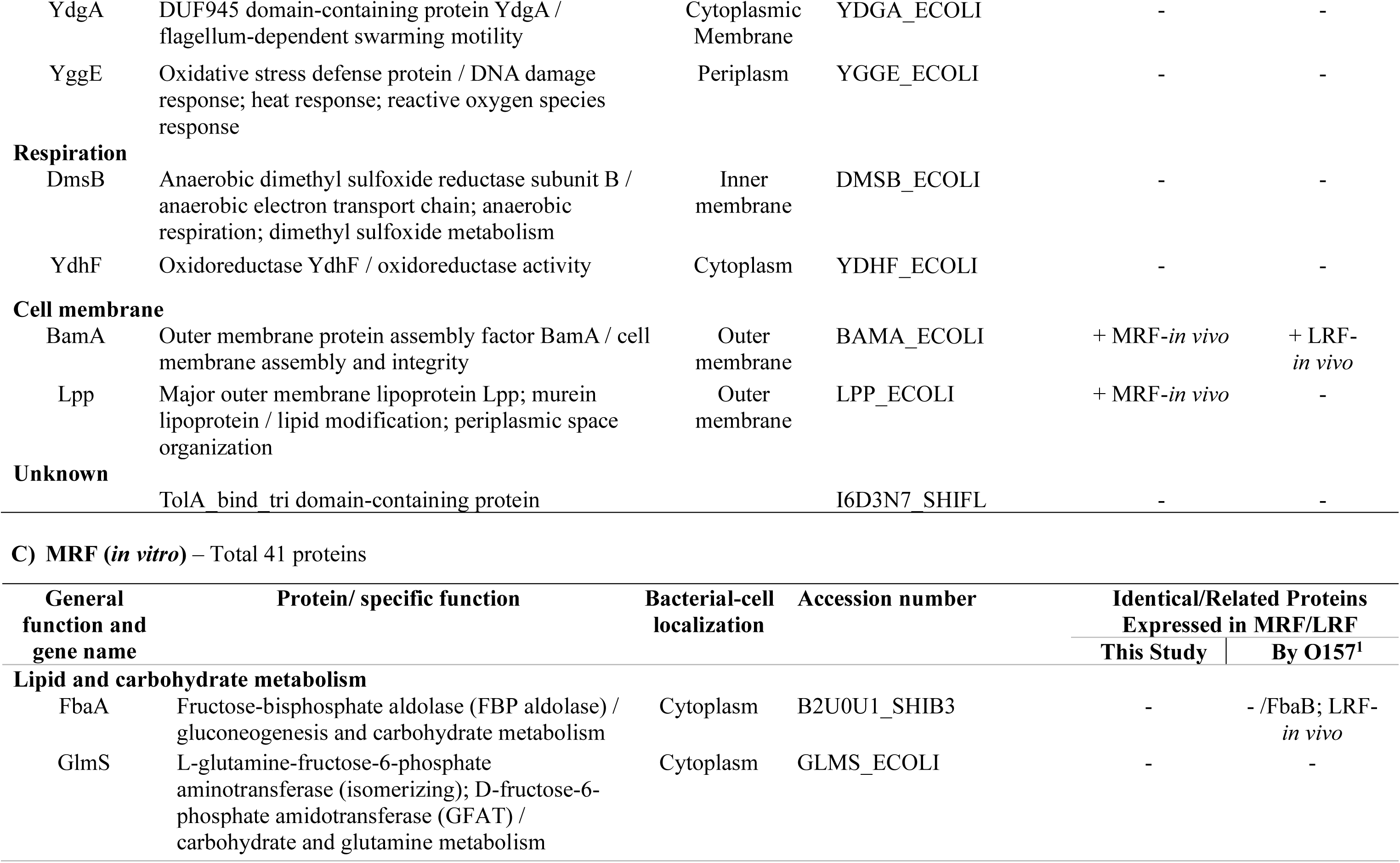
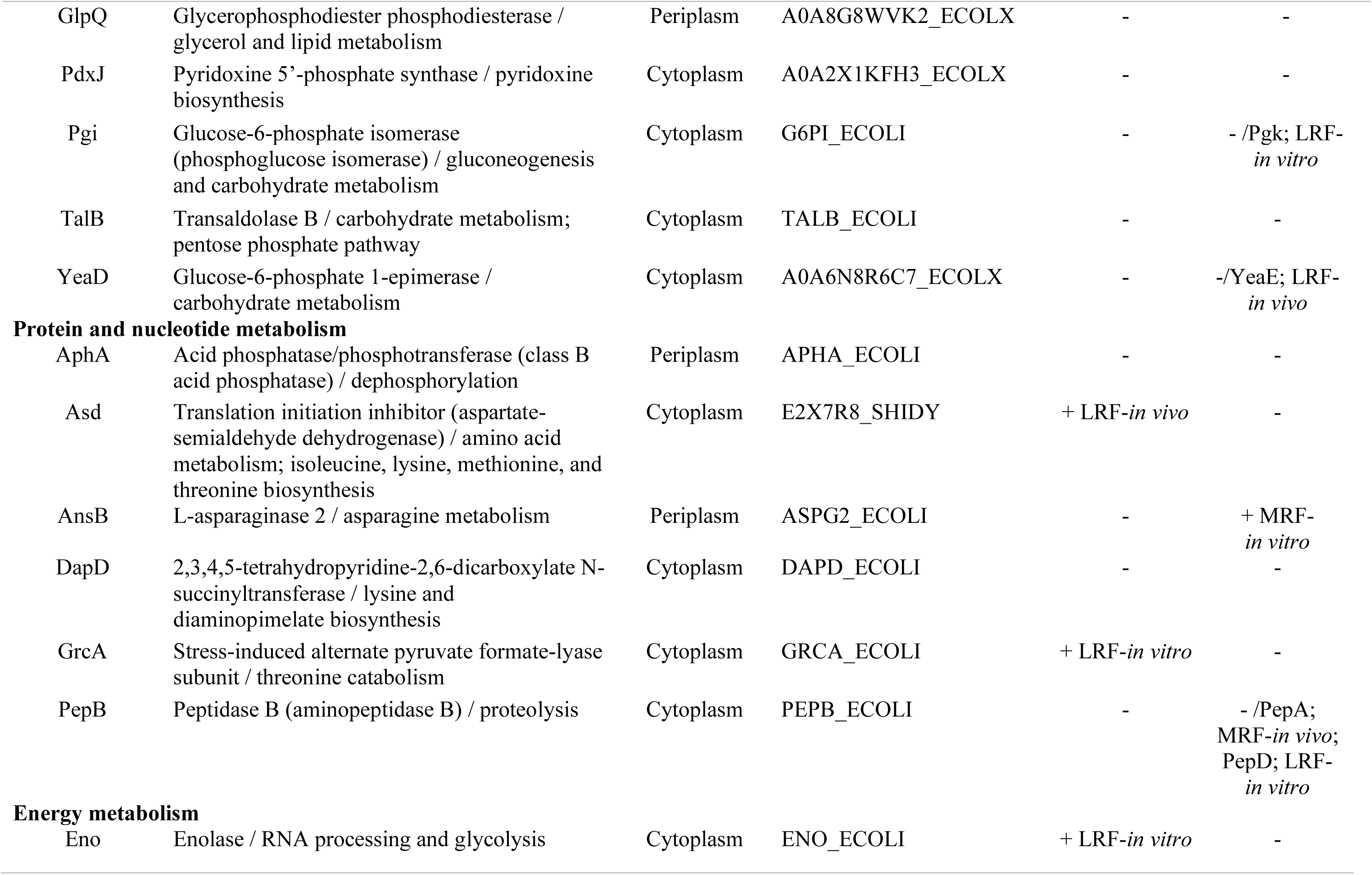
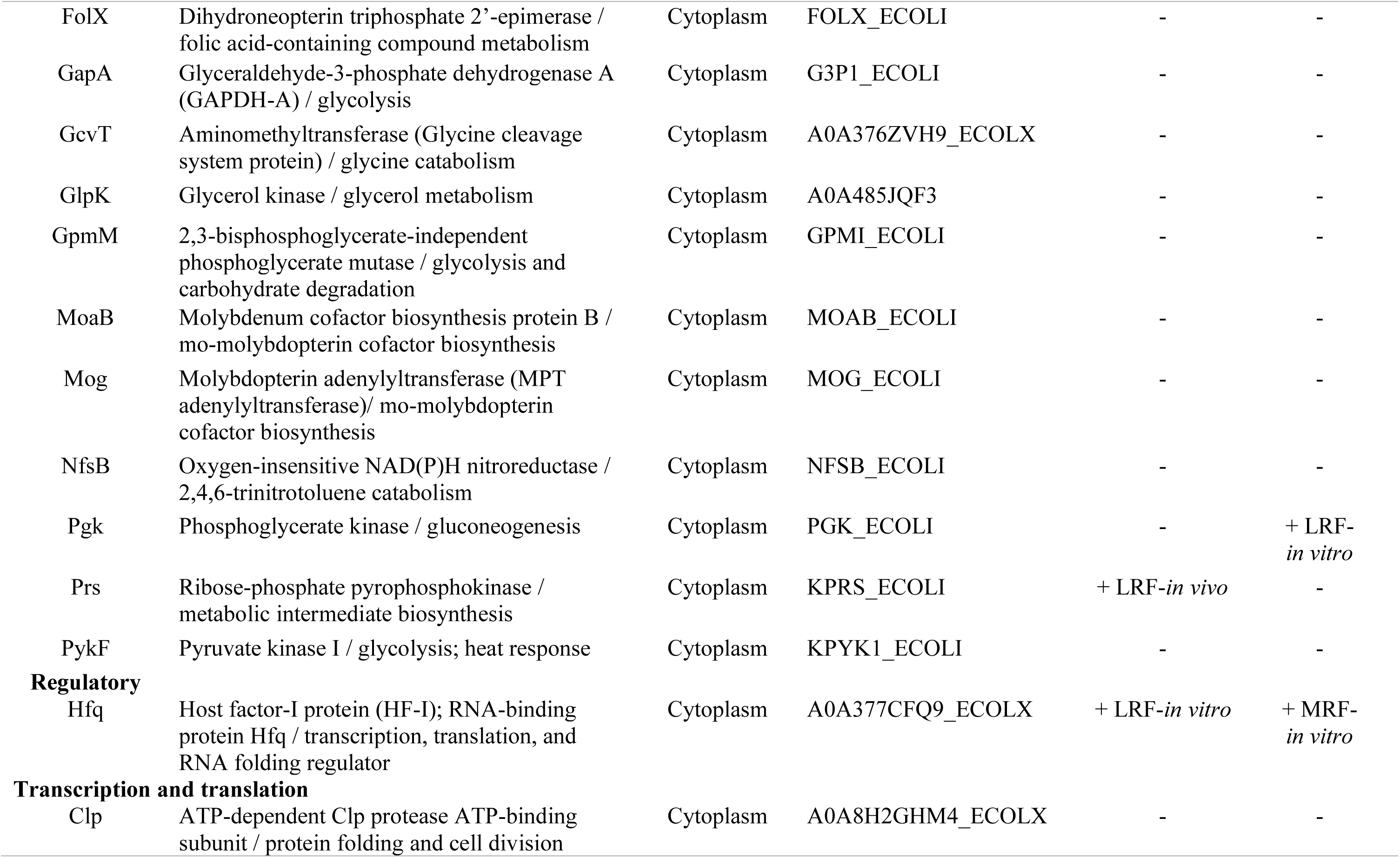
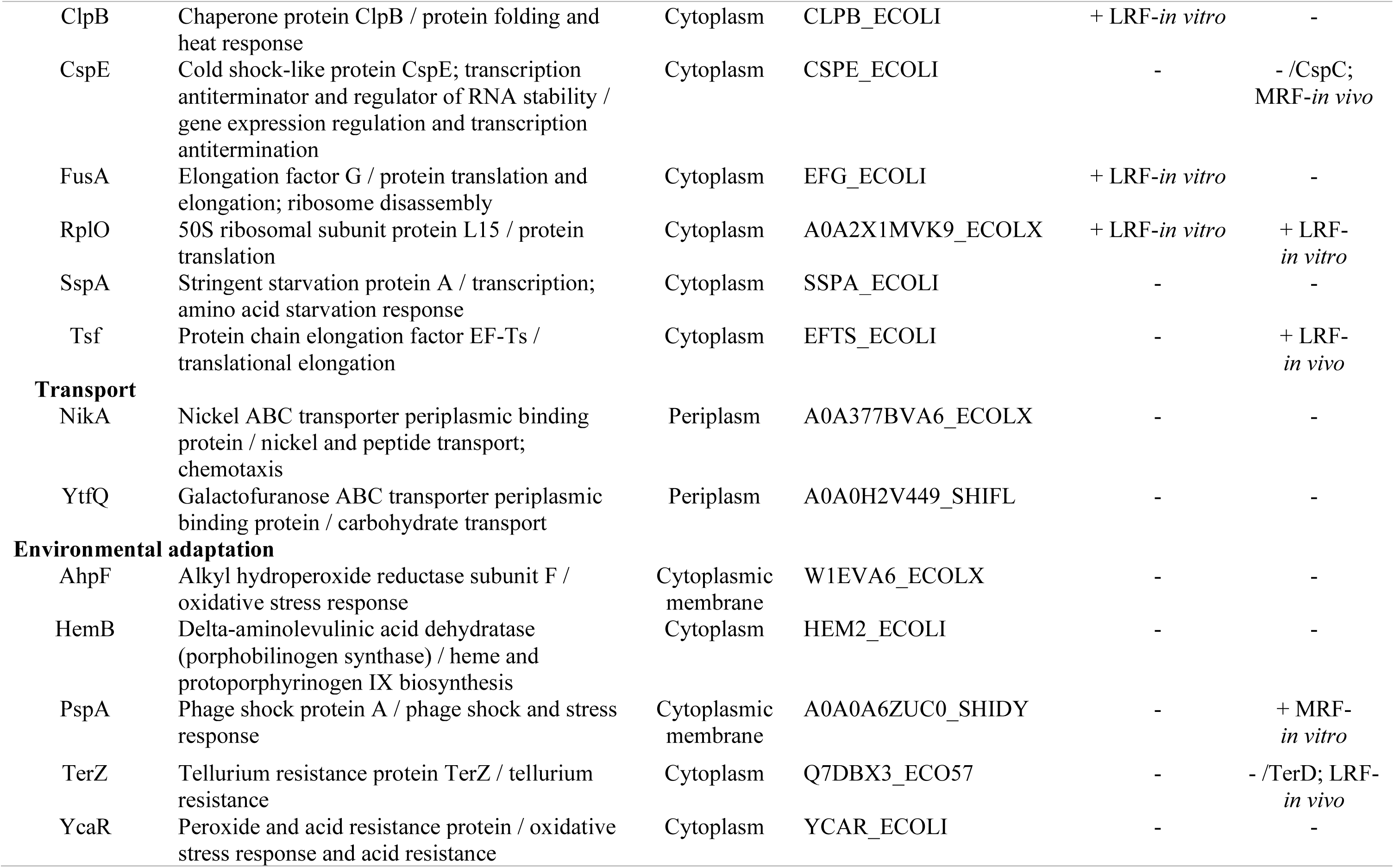
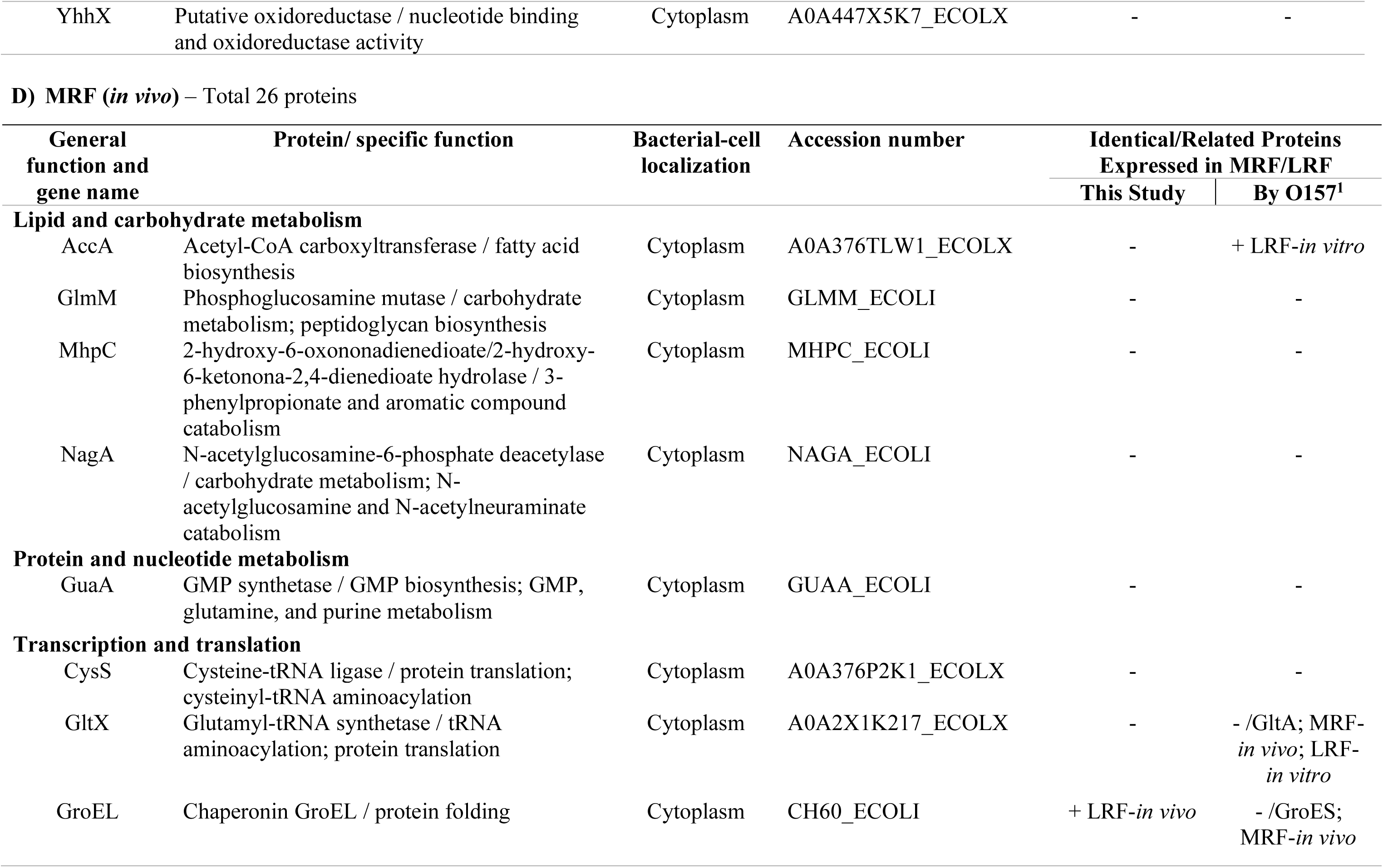
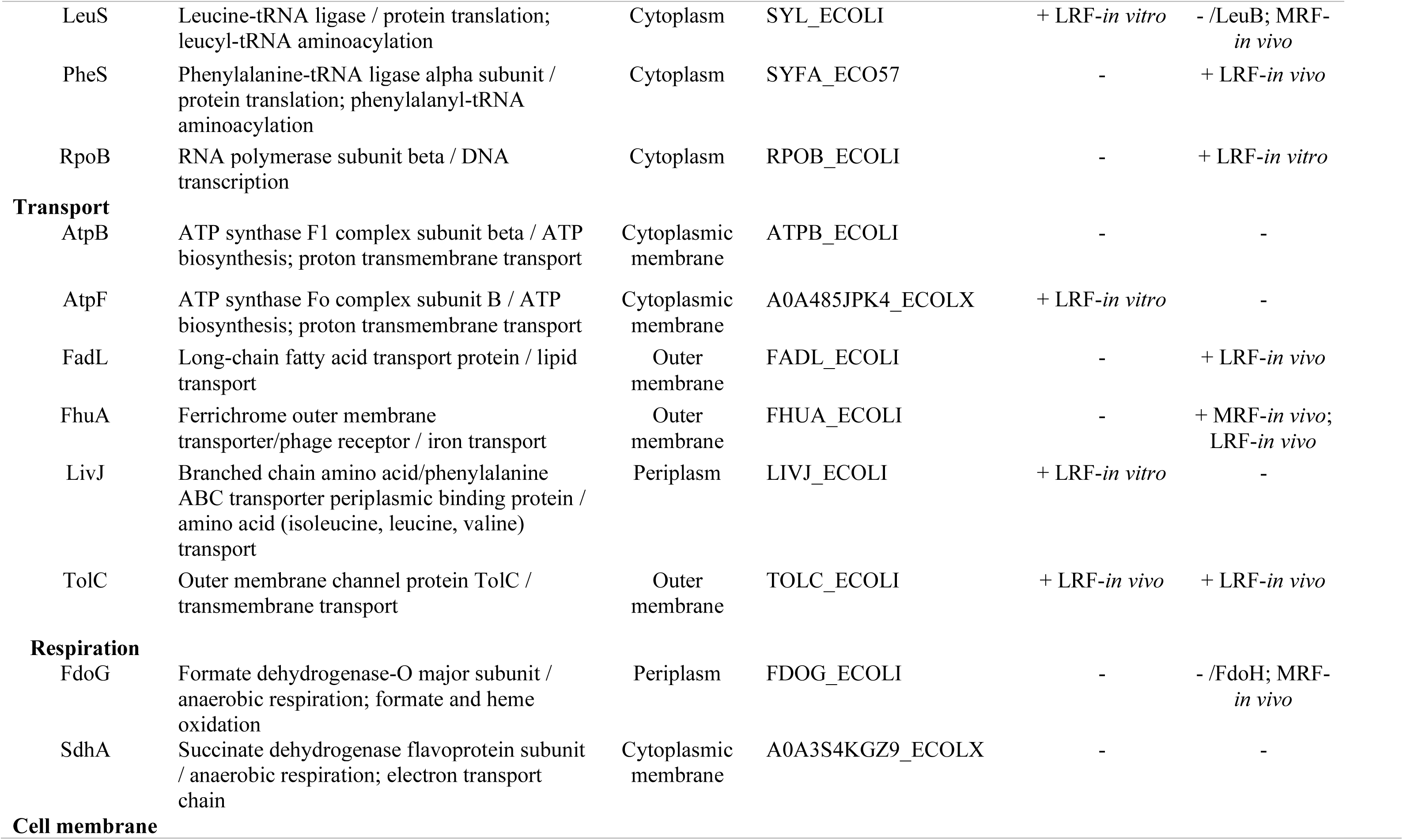
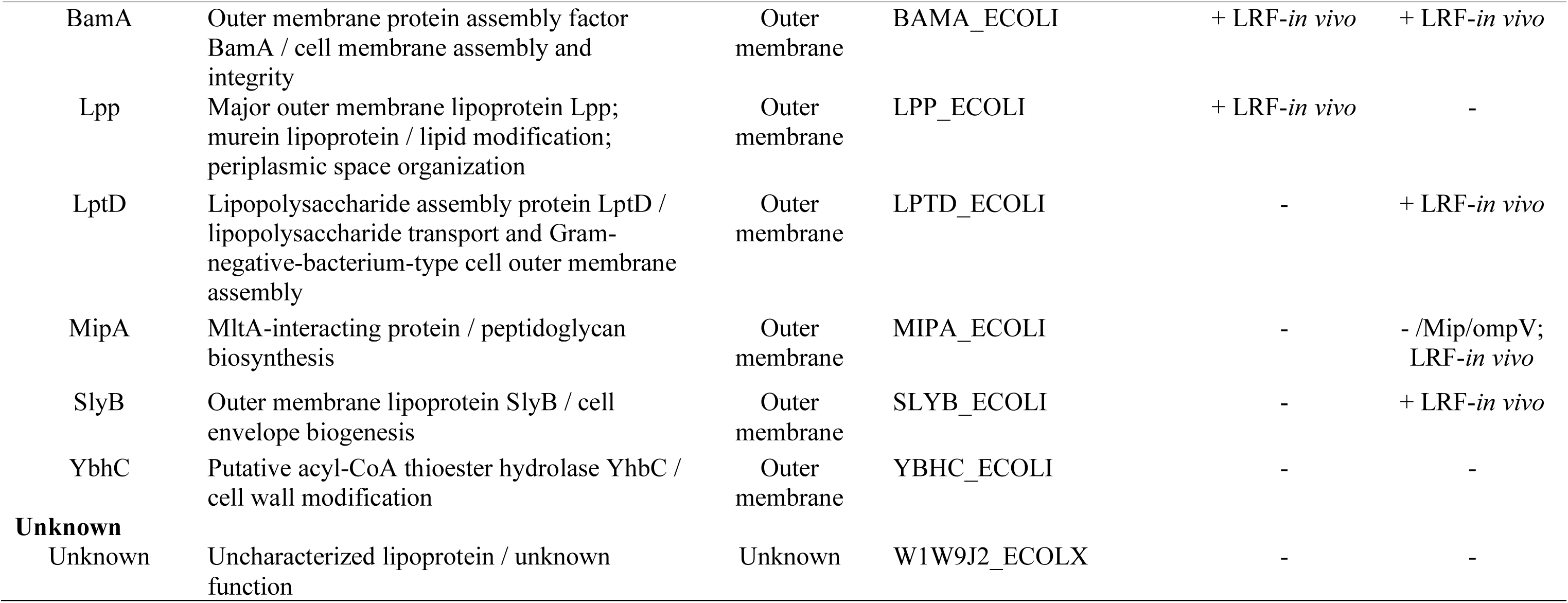
*E. coli* proteins differentially enriched in rumen fluid influenced by the maintenance (MRF) and lactation (LRF) diets, *in vitro* and *in vivo*, based on reference-free iTRAQ data analysis.

Few of the top 50 proteins with differential expression may have contributed to the outlier serotype-proteomes observed in the multivariate analysis (Figs. S4-S5; Tables S15 -S16). For instance, in LRF, proteins that were expressed by STEC O26:H11 and STEC O111:H8 *in vitro* (OmpA, OmpT, OmpX, NmpC, OmpC, FecA) were expressed *in vivo* by serotype O145:NM (Fig. S5). Similarly, some proteins were expressed *in vitro* by serotype O145:NM alone (UbiD_1, DcyD, HupB, RplL, NapA, NanA, RplC); these differences may have resulted in the serotype O145:NM being an outlier *in vitro* in LRF (Figs. S4 and S5). On the other hand, STEC O111:H8 was an outlier in LRF-*in vivo*, as it did not express RpS6, YjbH proteins under this growth condition (Figs. S4 and S5). In MRF the only outlier was the serotype O145:NM, under the *in vitro* condition (Fig. S4), expressing proteins otherwise expressed by the other two serotypes *in vitro* under *in vivo* growth conditions (ArgT, DcyD, NanA, SucC, YbiB, FumA, GldA_1, XdhD, AspA, EspC, UbiD; Fig. S5). While serotype O145:NM being an outlier under *in vitro* conditions in LRF and MRF may be associated to its porcine origin or lack of toxin genes, no such differences were observed *in vivo* ruling out this possibility.

## DISCUSSION

Besides the association with contaminated meat, salad, ice, and lettuce related outbreaks as discussed above (*see Introduction*; [9–12, 14–21, 23–25], STEC serogroups O26, O111 and O145 have also been recovered from various cattle samples using improved detection methods [55, 56]. For instance, multiplex PCR and immunomagnetic separation (IMS) enabled consistent isolation of STEC serogroups O26, O121, and O103 from cattle feces on a feedlot tested [56].

STEC O26 was also the second-most common serogroup to be isolated, using similar methods, from the feces of cattle in a Central US commercial feedlot with several isolates not carrying the *stx* genes [57]. Following sampling of hides and carcasses of 576 cattle, STEC serogroup O145 was determined to be second-most prevalent after STEC O157 on cattle hides using culture, PCR, and mass spectral analysis [58]. In another study, selective culture and PCR evaluation of recto-anal junction samples from 200 steers yielded STEC serogroups O26, O157, O145, O103 and O121 [59]. Globally, using different selective culture methods non-O157 STEC serogroups including O26, O103, O145, O121, have been isolated from food-animals consistently [60–63]. Interestingly, with the exception of one report indicating absence of toxin genes in the isolated non-O157 STEC[57], majority of the studies showed various combinations of the *stx_1_*, *stx_2_*, *eae*, and *ehxA* genes in the serogroups isolated from cattle [58, 61, 62, 64, 65]. Additionally, a variety of subtypes of *stx_1_* and *stx_2_* Shiga-toxins were identified in non-O157 STEC isolated from cattle feces [66].

Although there have been increased isolation of non-O157 STEC from cattle, little is known on the adaptative responses of these bacteria while in the rumen which is the first, largest and most complex compartment of the bovine stomach [30, 39, 67–71]. STEC need to survive the diverse microbiota, and their metabolic byproducts influenced by the diet in the rumen, in order to subsequently colonize the bovine intestinal tract [30, 39, 67–72]. In this study, we demonstrated that, as reported previously for STEC O157, the non-O157 STEC growth is suppressed by LRF and MRF resulting in reduction in viable counts [28]. However, the three non-O157 serotypes had survival patterns similar to each other in both LRF and MRF, irrespective of the growth condition and the various combinations of the *stx* and *eae* virulence genes. In addition, survival patterns were similar to the control non-STEC *E. coli* Nal^R^ during *in vivo* growth (Figs. 3B and 4B; Tables S1 and S2). A greater reduction in viable counts was observed *in vitro* in LRF than *in vivo* (Figs. 3 and 4 ; Tables S1 and S2), especially with STEC O26:H11; the acidic pH and increased volatile fatty acid concentrations may have influenced this suppression (Fig. 1: Table 1A). Smith *et al* reported a similar observation in their *in vitro* studies where STEC O26 was the most susceptible to low pH and oxidative stress [73]. In this study, the LRF pH was acidic with a total VFA concentration of 141 - 230 μM/ml, where the higher VFA values were observed only *in vitro* in LRF (Fig. 1). In contrast, MRF had a close to neutral pH and lower VFA concentrations (Fig. 2) with minimal increase *in vitro*. The *in vitro* changes in total VFA concentrations post-incubation, especially in LRF, may have been due to anaerobic fermentation by commensals in the rumen fluid rather than the test bacteria; accumulation of various VFA *in vitro* over time in the absence of active rumination has been reported [28, 29, 68, 71]. Taken together these survival patterns and recorded VFA-levels clearly supports our previous observation that the *in vitro* conditions cannot mimic the dynamic rumen environment of a live animal [28] and shows that the presence or absence of the *stx* and *eae* genes has no impact on STEC survival in the bovine rumen.

Unlike our previous observations with STEC serotype O157 [28], the serotypes O26:H11, O111:H8 and O145:NM shared survival patterns with the control non-STEC *E. coli* Nal^R^ (Figs. 3 and 4) in this study which seemed to indicate a greater degree of relatedness of non-O157 STEC to commensal *E. coli* compared to STEC O157 [74, 75]. While the evolution of non-O157 STEC is not fully understood, each of its serogroup appears to have followed the trajectory of genetic acquisition of various genes including Shiga toxins and intimin by commensal or other non-commensal *E. coli* supporting the relatedness [76–79]. In the process, multiple lineages of STEC O26:H11 evolved over time with most strains producing only Stx1; the *stx*_2_ genes in this serogroup are a relatively recent acquisition [80, 81]. *E. coli* O111:H11, comprising several multi-locus enzyme electrophoretic types, derived from multiple diarrheagenic *E. coli*, is genetically more diverse than rest of the STEC O111 serogroup [82, 83]. Additionally, STEC O26 and O111 serogroups were also determined to be closely related to *E. coli* associated with urinary tract infections and meningitis [79].

Analysis of the proteomes expressed by the three non-O157 serotypes in the rumen/rumen fluid showed proteins specific for survival being predominantly expressed over virulence proteins as previously reported for O157 strains (Table 3; [28, 29]). The exceptions were, intimin involved in STEC adherence detected in LRF [53] and glutamine/arginine transport proteins associated with the acid resistance detected in LRF and MRF [54] using iBAQ (Table S3- S6). General protein functions included: metabolism (lipid and carbohydrate, protein and nucleotide, energy), regulation, translation and transcription, transport, environmental adaptation, respiration and cell membrane synthesis (Table 3). Compared to the *in vitro* proteomes, the non-O157 *in vivo* proteomes were exclusive to that growth condition with only 5 proteins (Asd, Prs, LeuS, AtpF, LivJ) being expressed both *in vivo* and *in vitro* (Table 3).

Proteins expressed *in vivo* irrespective of diet type were associated with protein folding (GroEL), transport (TolC), or cell membrane assembly/modification (BamA, Lpp) functions (Table 3); identical or related proteins were also identified in the *in vivo* O157 proteomes[28]. Additional proteins that overlapped between the O157 and non-O157 *in vivo*-LRF/MRF proteomes included those involved in translation (RpsB, RpsJ, RpsO, TrpS, Glt, Leu, PheS), transport (ArgT, FadL, FhuA), respiration (Fdo), and cell membrane assembly (LptD, Mip, SlyB) (Table 3; [28].

Overall, proteins involved in transport, adaptation, respiration and cell membrane biosynthesis were enriched *in vivo* while those associated with the metabolic, regulatory, and translational pathways were enriched under *in vitro* growth conditions (Table 3) further corroborating the disparate *in vitro* and *in vivo* growth conditions. Subtle differences between proteomes of the serotypes tested were more evident in LRF than MRF (Fig. S4 and S5), driven by the animal’s diet and growth conditions. However, overall, the serotypes expressed more similar proteins despite differences in origin (human and porcine) (Figs. S4 and S5).

This is the first study outlining the survival and protein expression patterns for serotypes O26:H11, O111:H8, O145:NM *in vivo* in the bovine rumen using the non-terminal animal model. Data presented here-in was derived from only one serotype/strain per serogroup and hence is not a comprehensive representation for the entire serogroup. However, a sampling of the rumen survival and adaptation for the non-O157 serogroups O26, O111 and O145 is provided. This data may be used as a baseline for future studies evaluating additional serotypes/strains within these non-O157 STEC serogroups. Since there is scope for developing/improving preharvest strategies, these explorative insights could also provide additional common/unique targets/conditions for STEC control. The proteins expressed *in vivo*, while being driven by the animal’s diet and the growth condition was also influenced by the relatedness of the serogroups, to each other and to commensal *E. coli*. The differences and commonalities in STEC O157 and non-O157 serogroup adaptations need to be factored in when designing optimal targeted or broad STEC control modalities, be it dietary/prebiotic, probiotic or with inhibitory molecules [27, 84–88].

## ETHICS STATEMENT

The animal study was reviewed and approved by the USDA-ARS-Institutional Animal Care and Use Committee, National Animal Disease Center, Ames, IA, United States.

## AUTHOR CONTRIBUTIONS

ITK conceptualized, designed and executed all the *in vitro* and *in vivo* experiments, interpreted culture and pH-VFA data, prepared samples for proteomics, analyzed and interpreted proteomics data by iTRAQ. JT determined VFA concentrations by gas chromatography, completed iBAQ and reference-free iTRAQ analysis and statistically analyzed resulting proteomics data output.

ENB categorized and tabulated differentially expressed proteins using NCBI tools and researched literature for manuscript. ITK, JT and ENB researched literature and drafted the manuscript. All authors reviewed, revised and approved the drafted manuscript.

## DATA AVAILABILTY STATEMENT

All relevant data are within the manuscript and its Supporting Information files.

## ACKNOWLEDGEMENTS

Skillful assistance with general husbandry and rumen sampling was provided by the NADC animal caretakers. Excellent technical assistance provided by *late* Mr. Bryan Wheeler with the collection and filtration of rumen fluid, and placement/harvesting of ruminal cartridges is sincerely acknowledged. iTRAQ proteomics was done at the Proteomics Division, ICBR, University of Florida, Gainesville, FL. We gratefully thank *late* Dr. John D. Lippolis at the Ruminant Disease and Immunology Research Unit, NADC for providing expert guidance in the use of MaxQuant for iBAQ-proteomics data analysis.

This research was supported by an appointment to the Agricultural Research Service (ARS) Research Participation Program administered by the Oak Ridge Institute for Science and Education (ORISE) through an interagency agreement between the U.S. Department of Energy (DOE) and the U.S. Department of Agriculture (USDA). ORISE is managed by ORAU under DOE contract number DE- SC0014664. All opinions expressed in this paper are the authors’ and do not necessarily reflect the policies and views of USDA, ARS, DOE, or ORAU/ORISE.

Mention of trade names or commercial products in this article is solely for the purpose of providing specific information and does not imply recommendation or endorsement by the U.S. Department of Agriculture. USDA is an equal opportunity provider and employer.

## SUPPORTING INFORMATION

### Supplementary Text Files

Supplementary File1_iBAQ-code.pdf

Supplementary File2_ReferenceFreeiTRAQ-code.pdf

### Supplementary Figures

**Figure S1:** Product labels for ‘Steakmaker’ and ‘Lactation Premix’ used in the animal diets.

**Figure S2:** Raw iBAQ and riBAQ normalized intensities for proteins in LRF (L) and MRF (M).

**Figure S3:** riBAQ: Histogram of proteins with log_2_ fold change values >1.

**Figure S4:** Multivariate similarities of proteomes expressed in LRF and MRF.

**Figure S5:** Top 50 proteins with most variation between the non-O157 serotypes in LRF and MRF.

### Supplementary Tables

**Table S1:** Recovery of bacteria from cartridges exposed to LRF in *in vitro* and *in vivo*.

**Table S2:** Recovery of bacteria from cartridges exposed to MRF in *in vitro* and *in vivo*.

**Table S3:** Bacterial proteins enriched in LRF; based on riBAQ values of L2FC > 1 only.

**Table S4:** Bacterial proteins enriched in MRF; based on riBAQ values of L2FC < -1 only.

**Table S5:** GO terms of bacterial proteins enriched in LRF.

**Table S6:** GO terms of bacterial proteins enriched in MRF.

**Table S7:** Centered log ratio normalized, iTRAQ list of bacterial proteins expressed *in vitro* in LRF.

**Table S8:** Centered log ratio normalized, iTRAQ list of bacterial proteins expressed *in vivo* in LRF.

**Table S9:** Centered log ratio normalized, iTRAQ list of bacterial proteins expressed *in vitro* in MRF.

**Table S10:** Centered log ratio normalized, iTRAQ list of bacterial proteins expressed *in vivo* in MRF.

**Table S11:** NOMAD normalized, iTRAQ list of bacterial proteins expressed *in vitro* in LRF.

**Table S12:** NOMAD normalized, iTRAQ list of bacterial proteins expressed *in vivo* in LRF.

**Table S13:** NOMAD normalized, iTRAQ list of bacterial proteins expressed *in vitro* in MRF.

**Table S14:** NOMAD normalized, iTRAQ list of bacterial proteins expressed *in vivo* in MRF.

**Table S15.** Bacterial proteins differentially expressed in LRF between the serotypes tested.

**Table S16.** Bacterial proteins differentially expressed in MRF between the serotypes tested.

